# The MyLo CRISPR-Cas9 Toolkit: A Markerless Yeast Localization and Overexpression CRISPR-Cas9 Toolkit

**DOI:** 10.1101/2021.12.15.472800

**Authors:** Björn D.M. Bean, Malcolm Whiteway, Vincent J.J. Martin

## Abstract

The genetic tractability of the yeast Saccharomyces cerevisiae has made it a key model organism for basic research and a target for metabolic engineering. To streamline the introduction of tagged genes and compartmental markers with powerful CRISPR-Cas9-based genome editing tools we constructed a Markerless Yeast Localization and Overexpression (MyLO) CRISPR-Cas9 toolkit with three components: (i) a set of optimized S. pyogenes Cas9-guide RNA (gRNA) expression vectors with five selectable markers and the option to either pre-clone or co-transform the gRNAs; (ii) vectors for the one-step construction of integration cassettes expressing an untagged or GFP/RFP/HA-tagged gene of interest at one of three levels, supporting localization and overexpression studies; and (iii) integration cassettes containing moderately expressed GFP- or RFP-tagged compartmental markers for colocalization experiments. These components allow rapid, high efficiency genomic integrations and modifications with only transient selection for the Cas9 vector, resulting in markerless transformations. To demonstrate the ease of use, we applied our complete set of compartmental markers to co-label all target subcellular compartments with GFP and RFP. Thus, the MyLO toolkit packages CRISPR-Cas9 technology into a flexible, optimized bundle to simplify yeast research.

## INTRODUCTION

The yeast *Saccharomyces cerevisiae* is a key model eukaryotic cell in part due to the ease of genetic manipulations resulting from its high rate of homologous recombination and the availability of stable plasmids. This led to the development of toolkits facilitating genetic manipulations, typically based on stable selectable markers (Sikorski and Hieter 1989; Longtine *et al.* 1998). However, the small set of available markers limited the number of manipulations possible without using methods to recycle markers based on counterselections (Alani *et al.* 1987; Storici and Resnick 2006) or the Cre-lox system (Jensen *et al.* 2014). The discovery of Clustered Regularly Interspaced Short Palindromic Repeats (CRISPR) and CRISPR-associated protein 9 (Cas9) has greatly improved the efficiency of genomic manipulations (DiCarlo *et al.* 2013; Doudna and Charpentier 2014). The Cas9 endonuclease can be directed to introduce a double stranded break (DSB) at a specific location determined by a guide RNA (gRNA). Subsequently, the cellular machinery attempts repair by either error-prone non-homologous end joining (NHEJ) or homology-directed repair (HDR). The increased propensity for HDR makes genomic edits based on homologous recombination over 1000-fold more efficient, alleviating the need to integrate stable selectable markers (Ryan *et al.* 2014).

In yeast, catalytically active *S. pyogenes* Cas9 has now been applied in multiple toolkits and contexts. Combined Cas9-gRNA expression plasmids were made, and gRNA cloning methods were simplified (Ryan *et al.* 2014; Laughery *et al.* 2015; Levi and Arava 2020). Groups also focused on multiplexing CRISPR, allowing multiple simultaneous integrations with methods to introduce multiple guides and donors (Ryan *et al.* 2014; Jakočiūnas *et al.* 2015; Mans *et al.* 2015; Horwitz *et al.* 2015; Jakočiūnas *et al.* 2015; Zhang *et al.* 2019). Others updated a toolkit based on Cre-LoxP recycling of selectable markers with CRISPR to make a markerless system for introducing untagged genes for metabolic engineering (Ronda *et al.* 2015; Jessop-Fabre *et al.* 2016). Further consolidation of guides and donor DNA has also enabled large-scale CRISPR-based screens (Sadhu *et al.* 2018; Bao *et al.* 2018; Guo *et al.* 2018).

While CRISPR-Cas9 systems have progressed, there have been limited efforts to establish a robust set of accompanying vectors for integrating compartmental markers and tagged or untagged genes of interest (GOIs) using Cas9. Thus, though CRISPR-Cas9 methodologies present advantages in terms of flexibility and efficiency, classic systems based on homologous recombination remain heavily utilized for protein tagging. Furthermore, plasmids are regularly used for the expression of fluorescent compartmental markers, leading to uneven expression within populations (Ryan *et al.* 2014).

Here we present an optimized CRISPR-Cas9 toolkit for the rapid introduction of tagged genes and compartmental markers. First, we simplified the Cas9/gRNA expression vector developed by Ryan *et al.* to enable PCR-free Golden Gate cloning of gRNAs into vectors with five different selections (Ryan *et al.* 2014). We then established novel “split selection” markers that improve transformation efficiencies by allowing transformation of linearized pCAS vectors with intramolecular recombination generating a selectable marker. To complement the pCAS vectors, we developed a set of Golden Gate cloning-compatible plasmids that can be used to rapidly create integration cassettes that can introduce a gene of interest (GOI) with three different N- or C-terminal tags at three expression levels. Importantly, these cassettes contain novel multi-purpose homology arms that allow gRNA-based targeting to seven different established safe harbor loci, helping ensure that an integration site is usually available. Finally, we generated a set of GFP or RFP-tagged compartmental markers to facilitate and standardize colocalization studies, aiming to impart the simplicity of plasmids to the stability of integration. Together, these optimized yeast pCas9 vectors and versatile integration cassettes form the Markerless Yeast Localization and Overexpression (MyLO) CRISPR-Cas9 toolkit.

## RESULTS

### Optimizing use of Cas9 expression vectors in *S. cerevisiae*

To create a set of flexible Cas9-gRNA expression cassettes we modified an established pCAS vector (Addgene #60847) (Ryan *et al.* 2014). This vector expresses *S. pyogenes* Cas9 and a gRNA comprised of the 5’-cleaving hepatitis delta virus (HDV) ribozyme, a 20 bp proto-spacer responsible for Cas9 targeting and a scaffold mediating Cas9 interactions. *S. pyogenes* Cas9 targets regions that match the protospacer sequence and are followed by the protospacer adjacent motif (PAM) - NGG-, cutting 3 bp upstream of the PAM (Jinek *et al.* 2012; Gasiunas *et al.* 2012). We first substituted the *E. coli*/yeast Kanamycin resistance (KanR) cassette for a Hygromycin resistance (HygR) cassette and, in each plasmid, replaced the protospacer with a cleanly excisable stuffer containing a NotI and two BsaI cut sites, respectively generating pBBK94 and pBBK95.

These vectors can be used by excising the stuffer and co-transforming the pCAS with donor DNA and a protospacer fragment made by dimerizing two primers **(Figure 1a).** Inside the cell homology directed repair (HDR) introduces the protospacer into the pCAS allowing expression of Cas9 and a complete gRNA. Selecting for pCAS is sufficient for identifying integrants, and once identified, the selection can be dropped allowing the cells to discard the plasmid resulting in a “markerless” genome edit. While flexible, this approach depends on error free recombination to generate the gRNA. Unfortunately, either non-homologous end joining (NHEJ) occurring at the guide site or errors in gRNA primers can result in failure to activate Cas9 and the production of non-modified colonies.

**FIGURE 1.**
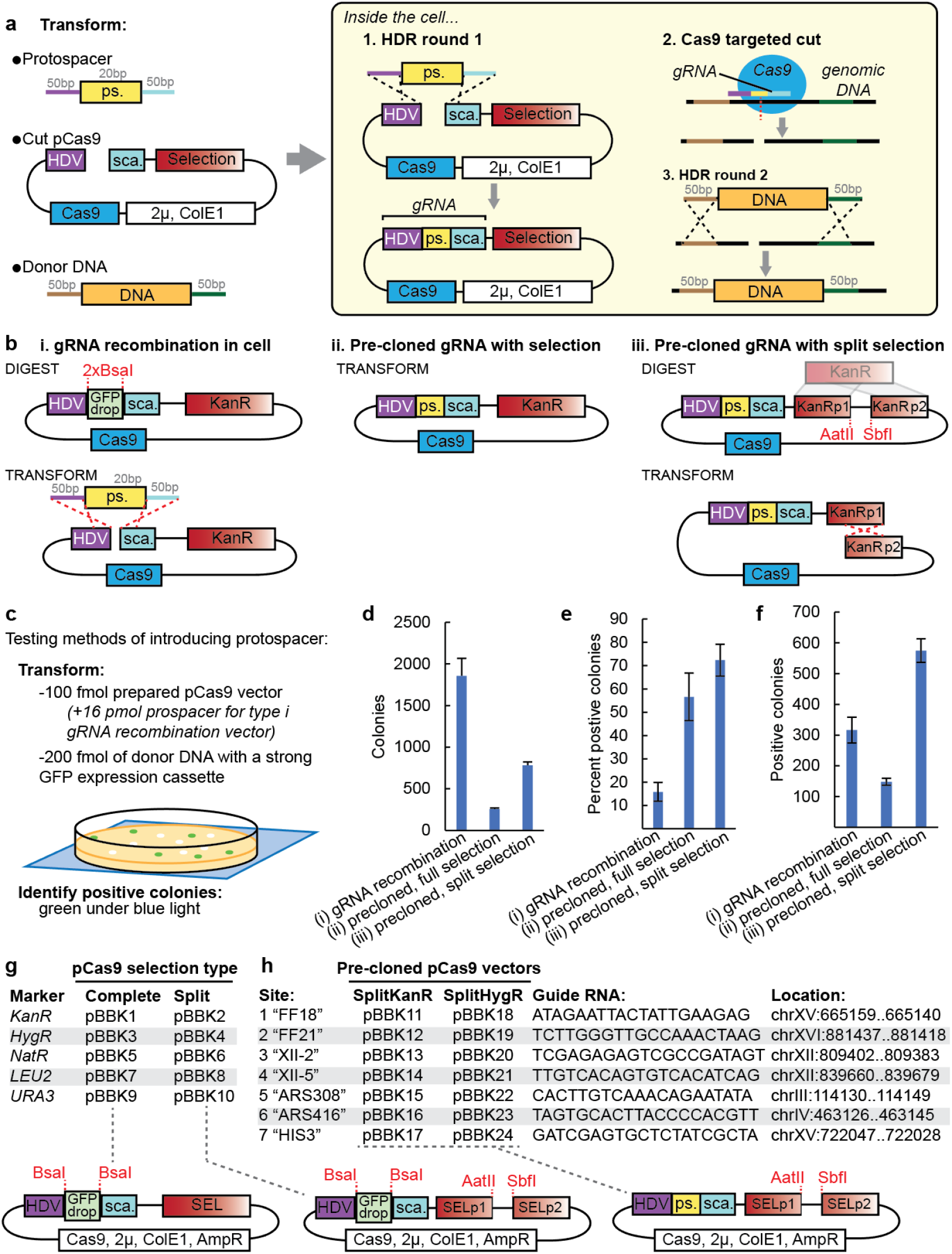
New Cas9 expression vectors for improved markerless CRISPR transformations. **(a)** Classic approach for integrating DNA in S. cerevisiae with CRISPR-Cas9. Three DNA fragments are cotransformed: a protospacer (ps.), a cut Cas9 vector and donor DNA. In the cell, [1] the pCAS vector is repaired, introducing the protospacer upstream of a scaffold (sca.), together creating a complete gRNA. [2] The gRNA directs Cas9 to cut at a site specified by the protospacer, which is then [3] repaired by the cell using the donor DNA. **(b)** Alternative strategies for introducing the gRNA and Cas9 beyond (i) the classic recombination approach include either pre-cloning the protospacer into a Cas9 vector with a (ii) complete selection or (iii) a “split selection” that must be repaired in the cell prior to expression. **(c)** To test the efficiency of the three approaches, 100 fmol of each Cas9 vector, with 16 pmol protospacer for the type i gRNA recombination vector, and 200 fmol of a strong GFP expression integration cassette (TDH3p-GFP) were transformed. Green colonies under a blue light indicated successful transformations. The resultant **(d)** total number of colonies, **(e)** percent of colonies that were positive and **(f)** total number of positive colonies were determined. **(g)** A series of Cas9 vectors with five types of complete or split selections. **(h)** A series of split selection Cas9 vectors with pre-cloned gRNAs for seven sites used in this toolkit. n = 3 and error bars indicate standard deviation.

Consequently, we investigated improving transformation efficiency by modifying how the protospacer is introduced. Initial tests of pre-cloned protospacers resulted in substantially lower transformation efficiencies, leading us to hypothesize that linearization of the pCAS is required for high efficiencies (data not shown). Therefore, we shifted the site of linearization to the selection cassette such that selection would depend on repair. We built a novel KanR-based “split selection” (SplitKanR) cassette by fusing the first two thirds of KanR to the final two thirds with a short linker containing restriction enzyme cut sites **(Figure 1b iii).** The original pCAS KanR marker was replaced with SplitKanR, and an ampicillin resistance cassette for *E. coli* selection was added. Transformations with this plasmid linearized at the SplitKanR should result in increased efficiency because there is more homology for HDR-based repair, the homology is intramolecular and the process selects for error free repair.

We compared the transformation efficiencies of the three gRNA introduction strategies: recombination at the gRNA, pre-cloning the gRNA with a complete selection, and pre-cloning the gRNA with a split selection **(Figure 1b/c).** To do so, we transformed a strong GFP expression cassette into the safe harbour site FF18 and identified positive colonies by their green fluorescence **(Figure 1h).** Transformation tests used 200 fmol of donor DNA (from pBBK93), 100 fmol of prepared pCas9 vector and, for the first strategy, a 160-fold excess of the protospacer. The gRNA recombination vector resulted in the most colonies, while the pre-cloned full and split selection vectors respectively yielded 14% and 42% as many colonies **(Figure 1d).** This is consistent with reports that linearization improves CRISPR-Cas9 transformations and suggests that either the protospacer excess or NHEJ-based repair at the protospacer site further increases the number of transformants (Horwitz *et al.* 2015; Guo *et al.* 2018). However, the protospacer recombination approach yielded only 16% positive colonies (green under blue light), roughly 4-fold less than either pre-cloned approach **(Figure 1e).** The colonies that were negative with the pre-cloned approaches were likely a result of genomic repair by NHEJ. Thus, the pre-cloned split selection pCAS yielded the most positive colonies, highlighting that this novel approach can take advantage of linearization-mediated transformation efficiency improvements while avoiding the generation of inactive gRNAs **(Figure 1f).**

We next generated and validated a series of pCAS vectors with a variety of complete and split selections for either single step gRNA recombination transformations or pre-cloning of gRNAs **(Figure 1g).** We included vectors for the gRNA recombination method as it is faster and easier for one-off transformations or efficient guides. To facilitate gRNA cloning, all vectors contain a *Bsa*I-flanked GFP dropout cassette at the protospacer site for PCR-free Golden Gate cloning (see Supplementary Information 2 for cloning guidelines). The *URA3* selectable marker in pBBK9 is also *BsmB*I-flanked to simplify introduction of alternate selections with Golden Gate cloning. We also cloned seven guides into both the SplitKanR and SplitHygR vectors **(Figure 1h).** These guides correspond to six commonly used safe harbor sites FF18/21 (Flagfeldt *et al.* 2009), XII-2/5 (Fabre *et al.* 2016), ARS308/416 (Reider Apel *et al.* 2017) and one that targets *HIS3* enabling screening. Collectively, our Cas9 vectors offer improved transformation efficiencies with flexibility in strategy and selections.

### A toolkit for rapid construction of integration cassettes

Classic approaches for expressing integrated tagged proteins often focus on introducing tags at the native focus of the gene of interest (GOI) (Fraczek *et al.* 2018). While this can be done markerlessly with CRISPR-Cas9, it requires the case-specific identification of a Cas9 cut site proximal to the 5’ or 3’UTR of the GOI and the amplification of donor DNA containing the tag with flanking homology specific to that site. Unfortunately, these cut sites can have lower efficiencies, and as the distance of the cut site from the gene increases the probability of the desired recombination event decreases (Supplementary Information 2). Furthermore, expression of heterologous genes requires an alternate approach, and colocalization studies often favor the ease of plasmid-based expression even though copy number variations cause significant variations in brightness (Ryan *et al.* 2014). CRISPR-Cas9 offers the alternative of easily introducing a complete expression cassette, containing the tagged GOI, at other loci such as the non-disruptive safe harbor sites **(Figure 1h).**

To facilitate this approach, we adopted a modular cloning scheme to generate plasmids that simplify building expression cassettes for untagged, GFP-, RFP- or HA-tagged GOIs (Lee *et al.* 2015) **(Figure 2a).** Each parent plasmid contains homology arms, a promoter, optionally a tag, a terminator and a *Bsa*I-flanked GFP dropout for introduction of a GOI by Golden Gate cloning. All tags contain glycine- and serine-rich linkers (Supplementary Information 1). Once assembled, *Not*I digestion linearizes complete integration cassettes for transformations. Parent plasmids are available featuring moderate to strong promoters (*RPL18Bp* < *TEF2p* < *TDH3p*) and with optional N- or C-terminal tags **(Figure 2b).** Both the GFP and the RFP are recently developed bright variants, ymNeonGreen and ymScarletI respectively (Botman *et al.* 2019). Microscopy and flow cytometry were used to verify ymNeonGreen was brighter than alternative GFP variants Envy and ZsGreenl (Supplementary Figure 1a/b).

**FIGURE 2.**
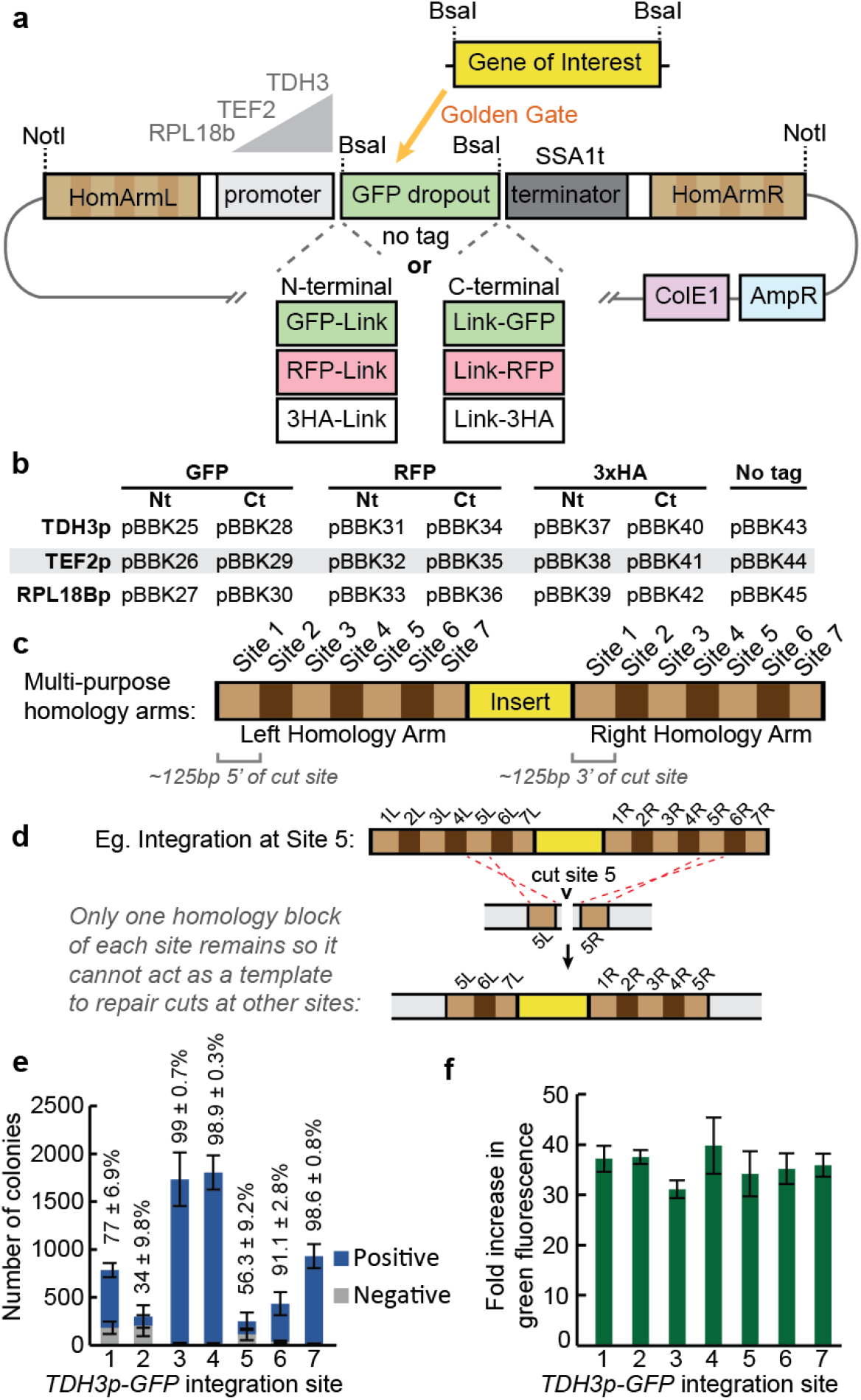
A set of markerless integration cassettes for introducing untagged, GFP-, RFP- or 3HA-tagged GOIs with multi-purpose targeting sequences. **(a)** Schematic of integration cassettes with left and right homology arms flanking a promoter at one of three strengths, a GFP dropout and a terminator. Some include N- or C-terminal GFP, RFP or 3HA tags as indicated. Golden Gate cloning with a BsaI-flanked gene of interest allows introduction of targets. **(b)** The integration cassettes parent vectors included in this kit. **(c)** The homology arms consist of roughly 125 bp regions either upstream (left arm) or downstream (right arm) of each Cas9 cut site described in Figure 1h, arranged in numerical order. **(d)** An example integration at Site 5 indicating that afterwards only one homology block of the other sites remains so this integrant will not interfere with subsequent transformations into those sites. **(e)** Transformation efficiencies at the different sites were determined by integrating a TDH3p-GFP cassette at each site and counting the total number of colonies as well as the number of colonies that were green under a blue light (positive). Transformations used 100 fmol of a pre-cloned split selection pCAS from Figure 1h and 200 fmol of the expression cassette. n = 4, twice each with SplitKanR and SplitHygR vectors. **(f)** To assess expression levels at the integration sites, TDH3p-GFP was introduced at each of site and fold induction of green fluorescence over wild type was measured by flow cytometry. n = 3; 10,000 cells/strain/replicate. Error bars indicate standard deviation.

These integration cassette parents feature unique multi-purpose homology arms **(Figure 2c).** Each arm contains a series of roughly 125bp sections homologous to regions upstream (left arm) and downstream (right arm) of the seven commonly used integration sites corresponding to the pre-cloned gRNAs in this kit **(Figures 1g, 2c).** The sequential orientation of Sites 1 to 7 on each arm means that once a cassette is integrated at one site it cannot act as a repair template for subsequent cuts at the other six sites **(Figure 2d).** This allows up to seven integration cassettes with the same homology arms to be sequentially introduced into a strain, facilitating gene dosage and colocalization experiments.

To validate this system, the KanR and HygR pre-cloned split selection pCAS vectors **(Figure 1g)** were used to integrate a strong GFP expression cassette (as in Figure 1c) into each integration site **(Figure 2e).** Transformations resulted in 250 to 1800 colonies, with integrations into Sites 3, 4 and 7 resulting in greater than 98% positive colonies. Integrations into Sites 2 and 5 yielded fewer than 60% positive colonies, which was unexpectedly poor relative to previous reports (Reider Apel *et al.* 2017; Bourgeois *et al.* 2018), though still sufficient for straightforward integration of cassettes. The drop in efficiency may reflect some structural feature of the homology arms or GFP expression cassette. The latter is likely for Site 2 as subsequent transformations into Site 2 for Figure 4c yielded 88% positive colonies by PCR (data not shown). Having confirmed the cassette could be integrated at all sites, expression levels between sites were compared. Expression levels, assessed by measuring green fluorescence by flow cytometry, were similar between sites at an average of 36-fold background fluorescence **(Figure 2f).** Together, the cassettes developed here can be integrated at seven gRNA-determined sites and result in expression levels uninfluenced by the integration site.

To confirm the functionality of all parent cassettes in Figure 1b, compartmental markers were cloned into each and then transformed into Site 1. For N-terminal tagging with fluorophores a plasma membrane marker comprised of two phospholipase Cδ Pleckstrin homology domains (2xPH) (Levine and Munro 2002) was used while the mitochondrial marker preCOX4 (Sesaki and Jensen 1999) was used for C-terminal tagging. When integrated, microscopy showed GFP-tagged **(Figure 3a/b)** and RFP-tagged **(Figure 3c/d)** versions were functional. Expression from tag-free cassettes was demonstrated by introducing GFP **(Figure 3e).** Hemagglutinin tagging cassettes were tested by introducing 2xPH and the vacuolar-localized protein PrcI (Huh *et al.* 2003) and Western blotting **(Figure 3f).** In all cases expression levels predictably corresponded to the promoter used, demonstrating the integration cassettes can be used effectively to integrate target GOIs at adjustable expression levels.

**FIGURE 3.**
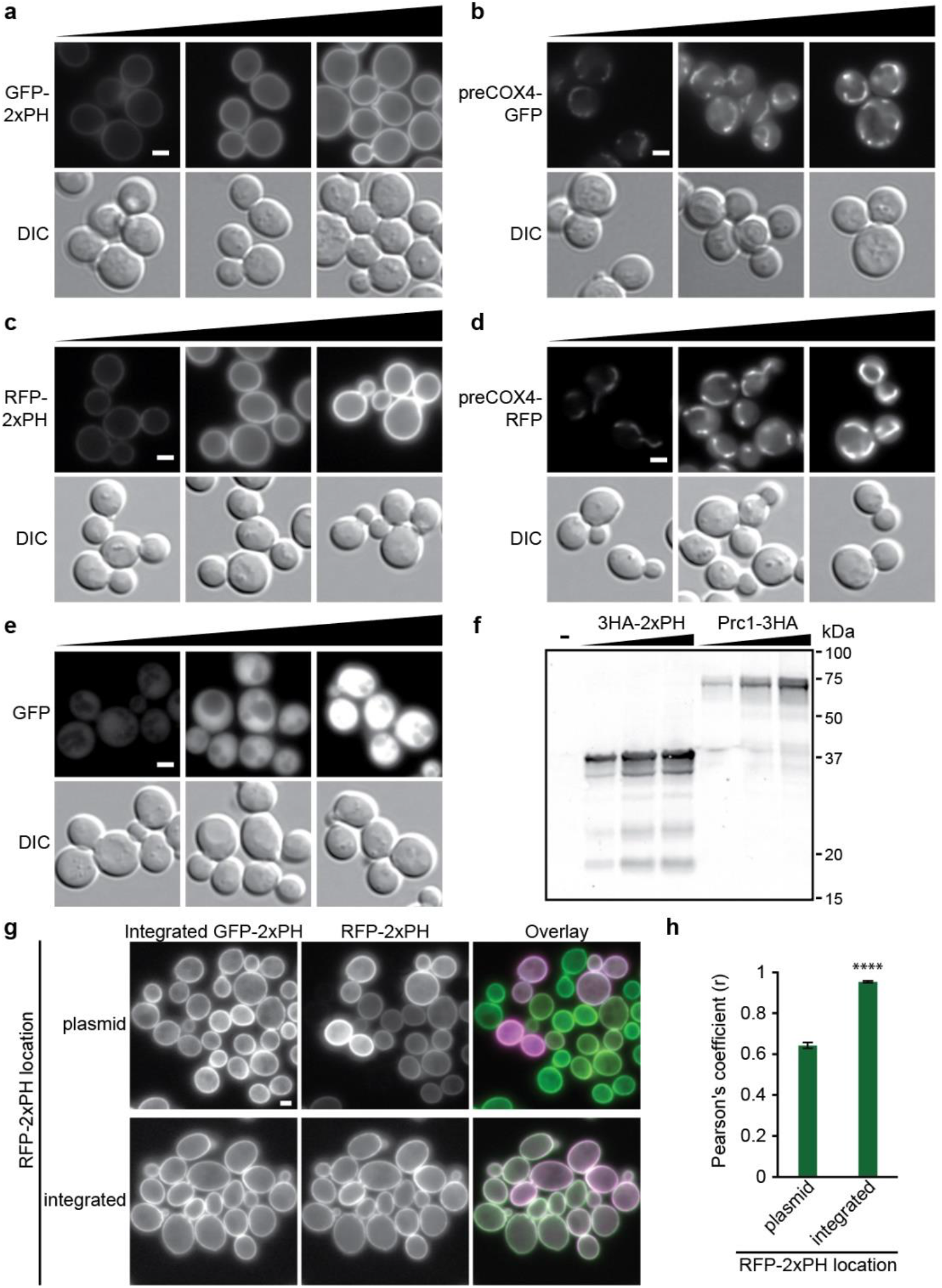
Validation of integration cassettes. All parent integration cassettes presented in Figure 2B were tested using compartmental markers. GFP-tagging cassettes were used to introduce **(a)** N-terminally tagged plasma membrane marker 2xPH or **(b)** C-terminally tagged mitochondrial preCox4 into Site 1 and the resultant strains were imaged. Black triangles indicate increasing promoter strengths; left to right: RPL18Bp, TEF2p, TDH3p. **(c)** RFP-tagged 2xPH and **(d)** preCox4 were also introduced at Site 1 and the strains were imaged. **(e)** GFP was introduced into untagged cassettes that were integrated at Site 1 prior to imaging. **(f)** HA-tagging cassettes were validated by introducing N-terminally 3HA-tagged 2XPH or C-terminally 3HA-tagged vacuolar marker Prc1 into Site 1 followed by an α-HA Western blot. n = 1. **(g)** Uniformity of expression was determined by comparing RFP-2xPH expressed from a low copy yeast centromere plasmid or after genomic integration in a strain with integrated GFP-2xPH. **(h)** Colocalization, assessed by Pearson’s coefficients, between GFP- and RFP-2xPH in the strains improved significantly when both markers were integrated. Two-tailed unequal variance t test. n = 6, >153 cells/condition/replicate. ****, P < 0.0001; error bars indicate SEM. Scale bars represent 2 μm.

To assess the evenness of expression from our integrated cassettes we introduced RFP-2xPH into our GFP-2xPH strain either on a low copy plasmid or into the genome at Site 2 **(Figure 3g).** As expected, there was substantially more variation in RFP-2xPH expression from the plasmid with some cells varying in fluorescence intensity or not expressing any detectable RFP-2xPH. This difference was quantifiable as a significant drop in the Pearson’s coefficient between the red and green channels when RFP-2xPH is expressed from a plasmid **(Figure 3h).**

### Integration cassettes with GFP- or RFP-tagged localization markers for colocalization studies

While fluorescently tagging a protein can reveal positional information, accurate identification of localization typically requires colocalization with a compartmental marker. To facilitate this type of work, we created a library of integration cassettes containing GFP- or RFP-tagged markers for 15 subcellular locations expressed from the *RPL18B* promoter (Figure 4a/b). The 21 markers selected were primarily well-characterized full length native proteins with localization confirmed in large-scale screens (Huh *et al.* 2003; Chong *et al.* 2015). Additionally, the transmembrane helix of Scs2 (Scs2tmh) was used to mark the ER (Loewen *et al.* 2007) and the 2xPH (Levine and Munro 2002) and preCox4 (Sesaki and Jensen 1999) markers were used for the plasma membrane and mitochondria respectively. Markers were C-terminally tagged except in cases where that had been shown to be disruptive, in which case they were N-terminally tagged (Chong *et al.* 2015) (Figure 4b asterisks).

**FIGURE 4.**
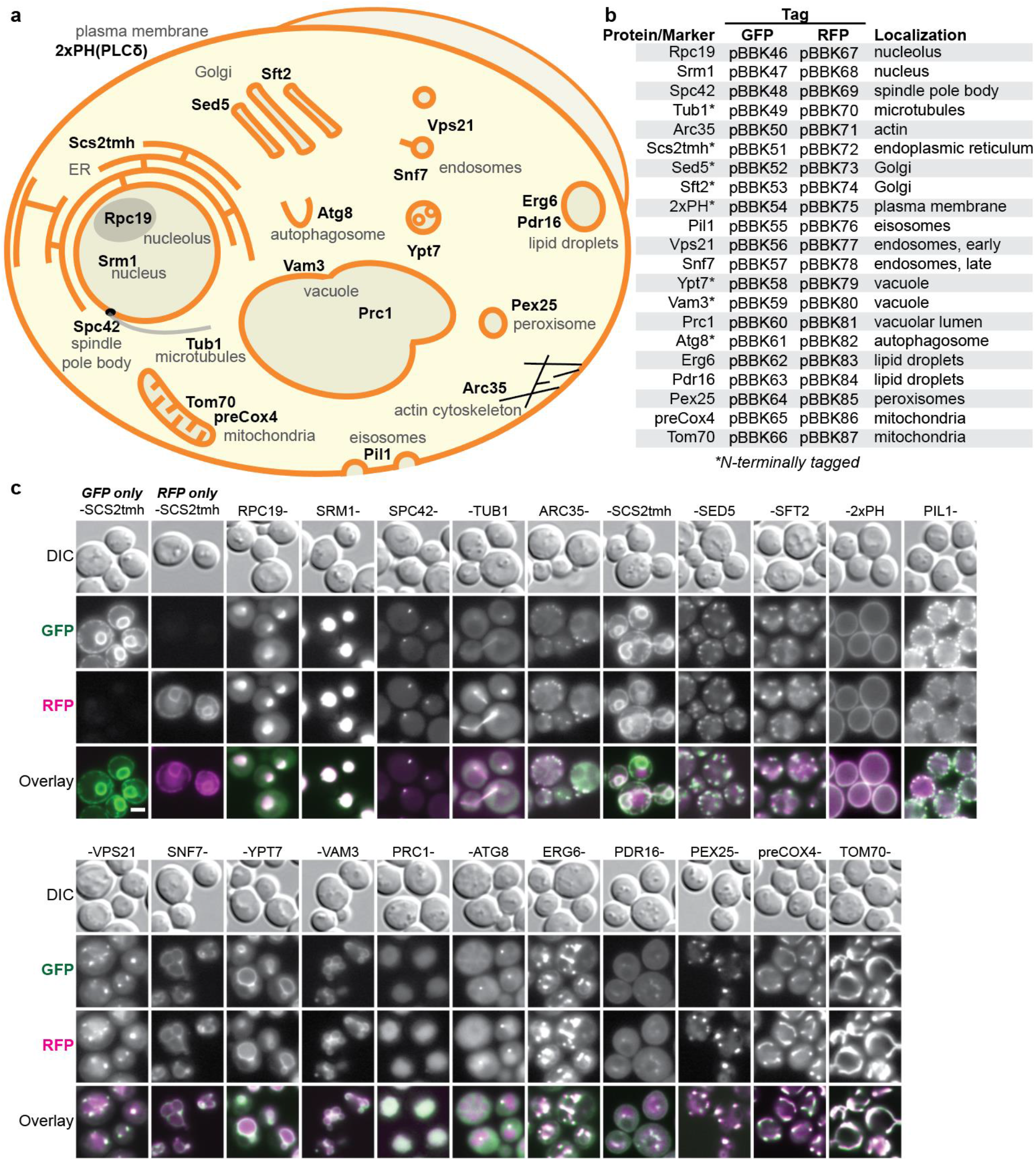
A library of integration-ready GFP- or RFP-tagged compartmental markers. **(a)** Schematic of a yeast cell with distinct structures and their associated markers indicated. Markers are complete yeast proteins except the transmembrane helix of Scs2 (Scs2tmh), a fusion of two phospholipase Cδ1 pleckstrin homology domains (2xPH(PLCδ)) and the Cox4 target peptide (preCox4). **(b)** Constructed integration cassettes expressing the indicated GFP- or RFP-tagged markers from RPL18Bp promoters. **(c)** The integration cassettes were introduced at Site 1 (GFP) and Site 2 (RFP) and colocalization was monitored by fluorescence microscopy. Scale bar represents 2 μm.

To validate the compartmental markers, we imaged strains with GFP- and RFP-tagged versions of each marker. The GFP and RFP cassettes were integrated at Sites 1 and 2 respectively and colocalization was assessed by fluorescence microscopy (**Figure 4c**). We observed both even expression levels and strong colocalization between GFP and RFP versions of each marker. Controls with only GFP-Scs2tmh or RFP-Scs2tmh demonstrated colocalization was not an artifact of bleed-through between channels. While there was some variation in marker intensities, consistent with variable protein stabilities, intensity was consistent enough to image all strains with the same exposure times (400ms GFP, 800ms RFP), thus streamlining microscopy. In some cases, the relative intensities of the markers varied with either the GFP (Pill) or RFP (Tubl, Arc35) markers being brighter, suggesting that in some cases one of the tags is more destabilizing or disruptive. Together, these markers demonstrate the ease of integrating tagged genes with the MyLO toolkit and should serve as useful tools for future colocalization work.

### Applying the toolkit for overexpression

The toolkit facilitates overexpression experiments by allowing sequential integration of a single cassette. Given the even expression across integration sites **(Figure 2F),** the relationship between cassette number and expression level should be linear if the cellular machinery mediating expression is not limiting. To establish this relationship, we sequentially integrated seven copies of GFP expression cassettes driven by *RPL18B, TEF2* and *TDH3* promoters at sites one through seven. Measuring the fluorescence of all strains, including intermediates, confirmed a linear relationship with copy number for each of the promoters used **(Figure 5a).** The strains with *TDH3p* and *TEF2p* cassettes were downward deflected and fit better with negative second order polynomials, indicative of expression limitation due to factors such as transcription factor titration or stress on protein biosynthetic machinery. This experiment also showed *TEF2p* is 5-fold and *TDH3p* is 11-fold stronger than *RPL18B.*

**FIGURE 5.**
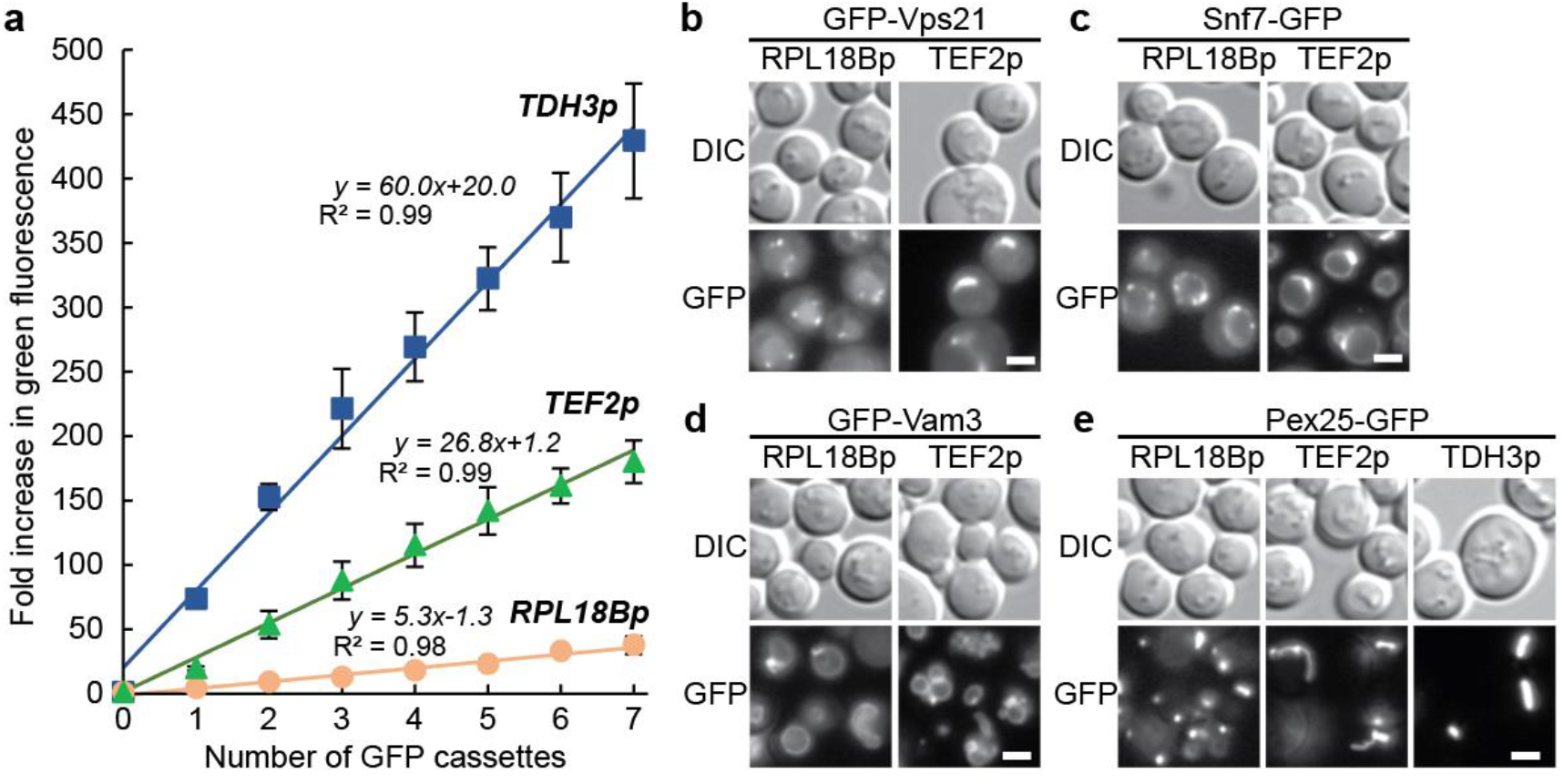
Using integration cassettes for overexpression. (**a)** Stepwise increases in GFP expression from cassettes with RPL18Bp, TEF2p and TDH3p promoters sequentially introduced at Sites 1 thorugh 7 was monitored by measuring green fluorescence with a flow cytometer. *n* = 3, 10,000 cells/strain/replicate. Fluorescence microscopy of yeast with GFP-tagged markers **(b)** GFP-Vps1, **(c)** Snf7-GFP **(d)** GFP-Vam3 and **(e)** Pex25-GFP displaying changes in localization as promoter strength increases RPL18Bp < TEF2p < TDH3p. Exposure times were adjusted to compensate for differing expression levels and highlight localizations. *n* = 2. Scale bars indicate 2 μm.

While multiple copies can be useful for gene dosage experiments or metabolic engineering, localization can be disrupted by expressing a gene with a stronger promoter. We observed such overexpression-induced mislocalization with some compartmental markers, as all were initially *TEF2p*-expressed to reduce exposure times. For example, the early endosomal Rab5 GTPase GFP-Vps21 shifted from small puncta to larger peri-vacuolar bars when *TEF2p*-expressed **(Figure 5b).** Likewise, *TEF2p-*expressed Snf7-GFP, a component of the endosomal sorting complexes required for transport (ESCRT-III) complex, accumulated in larger vacuolar puncta and on the vacuolar rim **(Figure 5c).** In other cases, overexpression disrupted compartmental shape. *TEF2p-*expression of the tSNARE GFP-Vam3 led to modest vacuolar fragmentation **(Figure 5d).** Strikingly, *TEF2p* expression of the peroxin Pex25-GFP resulted in a shift to bars with elongated tubules that sometimes extended from mother to daughter cells **(Figure 5e).** Further increasing expression by switching to the *TDH3* promoter resulted in loss of the elongated tubules and formation of intense bars. In all cases equivalent RFP-tagged markers localized similarly (Supplementary Figure 1c/d/e/f). In contrast, other tagged proteins maintained wild type localization when overexpressed, a subset of which we include as bright markers for the plasma membrane, endoplasmic reticulum and cytosol (pBBK88-93). These experiments demonstrate that overexpression studies can be conducted using the included promoters and homology arms.

## DISCUSSION

Our work provides an expansive toolkit optimized for Cas9-mediated introduction of tagged genes and compartmental markers into the yeast genome. Our set of pCAS-gRNA vectors provides flexibility with its five selectable markers and the option of using common pre-cloned gRNAs or introducing new guides by either Golden Gate cloning or homologous recombination in yeast. These pCAS vectors are complemented by a series of integration cassette parent vectors, simplifying the cloning of untagged and GFP, RFP or HA-tagged genes into expression cassettes, each of which can be integrated at any of seven sites. Also included is a collection of integration vectors with GFP- or RFP-tagged compartmental markers for use in colocalization studies. Together, these tools streamline many markerless genomic manipulations in yeast (Figure 6), supporting the rapid construction of more complex strains for basic and applied yeast research.

**FIGURE 6.**
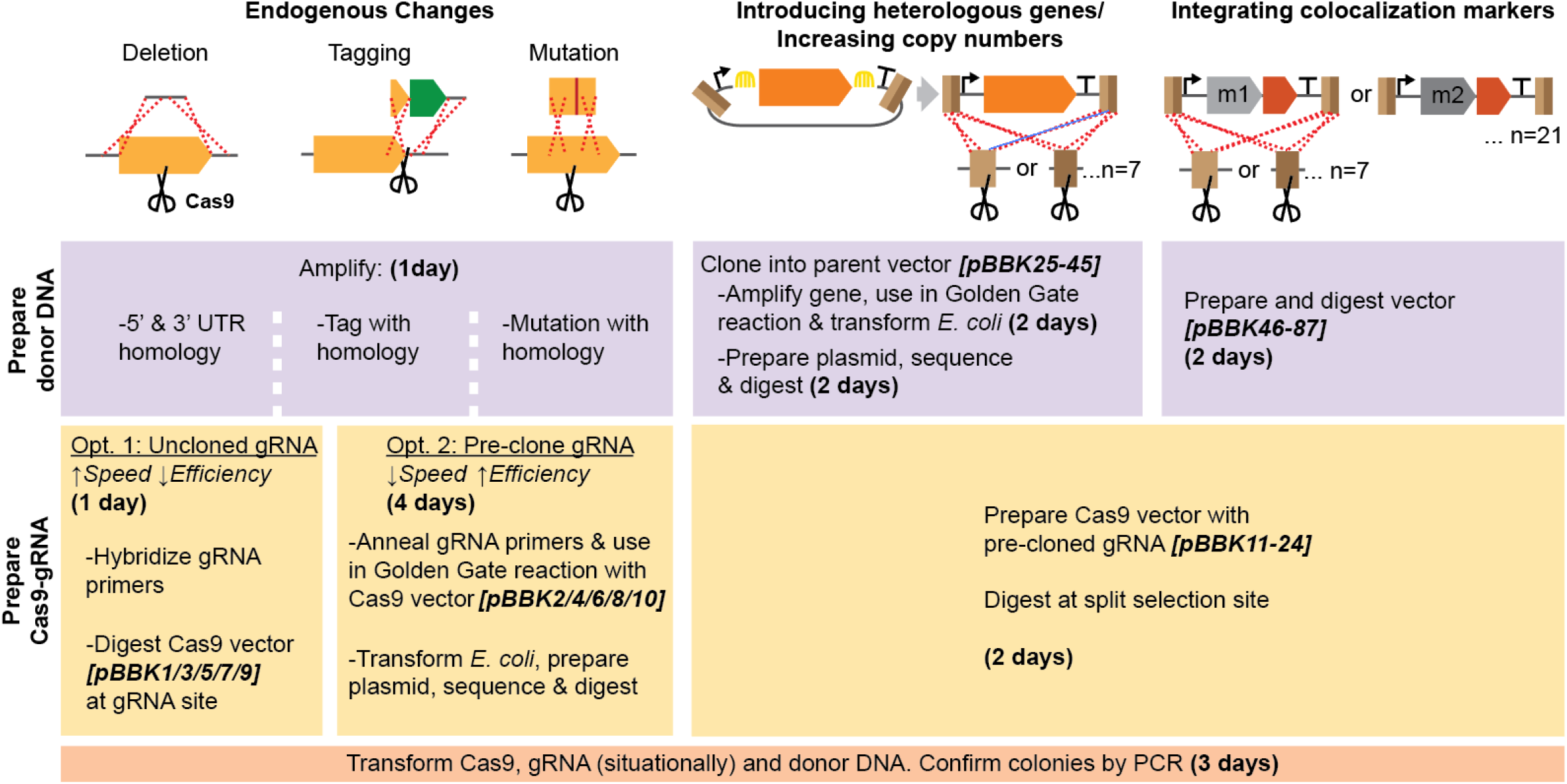
Overview of MyLO toolkit use cases and workflow. The toolkit facilitates both endogenous changes as introduction of heterologous genes. In all cases, donor DNA and Cas9 vectors require preparation (purple and yellow respectively), which can be done concurrently. Time required and vectors available are dependent on the method selected as indicated. In all cases without a cloning step, components can be prepared and transformed in a single day. Red dashed lines, crossover event; scissors, Cas9 cut; yellow arch, Golden Gate reaction; m, compartmental marker.

There is substantial variation in how the core requirements for Cas9-based transformations, namely the introduction of a constant Cas9 expression cassette with variable, yet matched, gRNAs and donor DNAs, is achieved. Some combine all elements on single plasmids (Bao *et al.* 2015; Vyas *et al.* 2018) whereas others rely on pre-loading a strain with either the Cas9 alone (DiCarlo *et al.* 2013; Fabre *et al.* 2016) or, with the use of an inducible Cas9 cassette, together with a gRNA (Degreif *et al.* 2018). While both approaches have been shown to work well, they either tend to require more complex cloning strategies or longer transformation protocols. In contrast, co-transforming a combined pCas9-gRNA vector with a donor, as we and others (Laughery *et al.* 2015; Generoso *et al.* 2016) have pursued, offers the simplicity of only needing to make small changes at a single site in the vector backbone to change the protospacer, with the convenience of a single step transformation into any strain. Given the utility of this approach we aimed to develop a vetted set of these vectors with a variety of markers, though we note that Addgene contains extensions to the Laughery *et al.* system and unpublished vectors from the Ellis lab (pWS158, 171-176) that fill similar niches. The availability of multiple markers is important for marker cycling, where dropping a selection after a transformation allows a plasmid to be lost fast enough that every third, or in some cases second, transformation can be done with the same selectable marker on the pCAS without needing a curing step. Lastly, we simplified this pCas9-gRNA strategy by making protospacer cloning optional, precloning common guides, and ensuring all cloning is Golden Gate compatible.

We found that linearization of the Cas9 vectors improved transformations (Figure 1). This, together with previous reports of gap repair increasing the percent of positive colonies (Horwitz *et al.* 2015), or both the number of colonies and the percent positive (Guo *et al.* 2018), could be explained by two mechanisms. First, as previously suggested, requiring gap repair of the Cas9 vector ensures that the cell is competent for homologous recombination, a prerequisite for efficient introduction of the donor DNA at the target site (Horwitz *et al.* 2015). Consistent with the expectation of a higher frequency of positive colonies, we observed a 28% increase in the percent of positive colonies with our split selection pCas9, relative to the circular vector. Second, the topology of linear DNA can promote improved transformation efficiencies, likely through increased DNA uptake (Raymond *et al.* 1999). Both this work, where linearized vectors resulted in 4-7x more colonies than transformations with circular pCas9, and that of Guo *et al.* support this mechanism as an important factor in improving Cas9-based transformation efficiency. Though our system uniquely implements an intramolecular gap repair strategy, the extent to which this boosts transformation efficiencies relative to intermolecular repair at either the marker (Horwitz *et al.* 2015; Guo *et al.* 2018) or to non-functional regions flanking the gRNA expression site (unpublished Ellis lab system; see pWS158 on Addgene) remains to be determined. Collectively, there is now strong evidence that gap repair improves CRISPR-Cas9 methods.

Our extended set of integration cassettes, ready for introducing a tagged or untagged GOI, provide a compromise between flexibility and simplicity. The use of a Golden Gate cloning strategy compatible with the modular cloning toolkit (Lee *et al.* 2015) should simplify the introduction of genes from existing vectors (see Supplementary Information 2 for cloning protocols). Furthermore, it should promote interchangability with a recent extension of the modular cloning toolkit capable of multiplexing and designed for more variable markerless CRISPR-based applications (Otto *et al.* 2021). In contrast to these alternative systems, the MyLO kit sacrifices some versatility by providing pre-built integration cassettes ready to introduce a GOI in a limited design space, three promoters with and without tags. This greatly simplifies cloning and minimizes sequencing requirements. Additionally, by avoiding multiplexing, we were able to introduce the unique multi-component homology arms that allow gRNA-guided targeting of a single cassette to any of seven sites. This allows many combinations of MyLO integration cassettes to be introduced both sequentially and repeatedly, without concern for clashing homology arms or the unavailability of a specific site, a frequent concern in metabolic engineering. Though crossover events could hypothetically occur between our integrated cassettes, in all cases where colonies were assayed by PCR we observed the expected phenotype, including after seven rounds of sequential transformations, suggesting the cassettes have high fidelity and stability (data not shown).

We used the integration cassette parents to make an extended set of GFP- and RFP-tagged compartmental markers designed to provide the even level of expression of integrations while approximating the ease of a plasmid transformation. Indeed, we found that expression levels were exceptionally uniform between cells with a given marker, but intensity did vary between markers expressed from the same promoter, likely due to different levels of recruitment to compartments and variations in protein stability. Given the comparable expression levels, it should be possible to build tester strains with multiple morphologically distinct compartments, such as the ER, endosomes and the plasma membrane, simultaneously tagged to rapidly identify the localization of a new protein. Our localization system does have the standard caveats associated with introducing second copies of genes: essential proteins inactivated by tagging will be possible to visualize though they may be mislocalized, and the untagged version of some proteins may partially outcompete the tagged version for recruitment to a given compartment. The ability to rapidly introduce a uniformly-expressed compartmental marker for colocalization should greatly facilitate localization studies and automated image analysis.

Initial overexpression of our compartmental markers inadvertently allowed us to interrogate the function of several yeast proteins. When overexpressed, Vps21 and Snf7 primarily accumulated in puncta on or adjacent to the vacuole. The localization of the endosomal Rab5 GTPase Vps21 to large peri-vacuolar puncta was similar to phenotypes resulting from constitutive activation or deletion of its GTPase activating protein (GAP) Msb3 (Lachmann *et al.* 2012; Nickerson *et al.* 2012). This suggests that there are sufficient levels of Rab5 guanine nucleotide exchange factors (GEFs) to activate and recruit overexpressed Vps21 to membranes, but not enough Msb3 to effectively terminate signaling, revealing a possible bias towards Rab5 activation (Cabrera and Ungermann 2013). Overexpression of C-terminal tagged Snf7 is known to disrupt ESCRT function leading to accumulation on endosomes and the creation of large vacuolar adjacent class E compartments (Raymond *et al.* 1992; Froissard *et al.* 2007; Bean *et al.* 2015). However, we also observed overexpressed Snf7-GFP on the vacuole, which supports the recent establishment of ESCRT function on the vacuole (Yang *et al.* 2021).

Another two markers, Vam3 and Pex25, led to larger abnormalities in compartmental morphology. Cells with overexpressed Vam3-GFP had fragmented vacuoles, a phenotype associated with loss of the tSNARE as it mediates homotypic fusion of vacuoles (Srivastava and Jones 1998; Lürick *et al.* 2015). Given that this function requires interactions with both the tether HOPS and other SNAREs, the overabundance of Vam3 likely disrupts vacuole fusion by upsetting the ratio of components forming this complex, reducing complete assembly (Alpadi *et al.* 2013; Lürick *et al.* 2015). Progressive increases in Pex25-GFP overexpression led to enlarged peroxisomes with elongated protrusions and then to intense bar-shaped structures. This is likely linked to the role of Pex25 in peroxisome biogenesis and regulation of peroxisome number and shape through interactions with the GTPase Rho1 (Marelli *et al.* 2004; Akşit and van der Klei 2018). Pex25 overexpression has been found to drive peroxisome proliferation and the emergence of clustered laminar membranes or juxtaposed elongated peroxisomes, which could be the bar-like structures we observed (Rottensteiner *et al.* 2003; Tam *et al.* 2003; Huber *et al.* 2012). The elongated tubules strikingly resemble those that form in the peroxisome-abundant yeast *Ogataea polymorpha* upon deletion of the dynamin Dnm1 that drives peroxisomal membrane scission, suggesting that overexpression of Pex25-GFP blocks the function of the yeast peroxisomal fission proteins Vps1 and Dnm1 (Smith *et al.* 2002; Nagotu *et al.* 2008; Akşit and van der Klei 2018). Together, these results highlight the rich information that can be obtained from overexpression experiments facilitated by this toolkit.

The MyLO toolkit provides a strong foundation for basic CRISPR-Cas9 genome editing in yeast, and could be expanded in numerous ways. These expansions could include adding new integration cassettes with multipurpose homology arms for different sets of sites, additional promoters including those allowing regulation of gene expression, and the inclusion of alternative compartmental markers. Likewise, the pCAS vectors could be modified to include new smaller or higher fidelity CAS proteins and should be easily adaptable to the method presented by Zhang *et al.* for multiplexing of gRNAs (Kim *et al.* 2017; Casini *et al.* 2018; Zhang *et al.* 2019; Xu *et al.* 2021). Indeed, the increased efficiency and simplicity of the split selection pCAS strategy could further improve current CRISPR multiplexing capabilities.

## MATERIALS AND METHODS

### Plasmids and Yeast strains

All plasmids used were from the MyLO Toolkit collection constructed here. They are listed in Table SI, and will be made available from Addgene. Plasmids were made using Golden Gate cloning or homologous recombination in yeast as outlined in detail in Supplementary Information 1 with primers listed in Table S2. Yeast strains, listed in Table S3, were made with CRISPR/Cas9-based transformations using MyLO Toolkit components. All plasmids were confirmed by sequencing and all integrants were confirmed by PCR.

### Fluorescence-based Yeast Transformation Assay

Assays of transformation efficiency were performed by integrating a strong GFP expression cassette (pBBK93, *TDH3p-Neon(gfp))* into wild type yeast using a lithium acetate-based transformation protocol (Gietz and Schiestl 2007). Briefly, 3 OD_600_ log phase yeast were transformed by incubating for 30 min at 30°C and then 42°C in a 150 μL final volume with concentrations as in Gietz *et al.* Twenty percent of the reaction was plated and incubated two days at 30°C prior to imaging on a Safe Imager 2.0 (Invitrogen) blue light source to identify positive, green colonies. In some cases, colonies were counted using the Fiji (Schindelin *et al.* 2012) Color Threshold tool followed by the Analyze Particles tool.

### Fluorescence Microscopy

Log phase yeast grown in synthetic selective media were imaged on slides using a DMi6000B microscope (Leica Microsystems) with an HCX PL APO 63x oil objective, an Orca R2 CCD camera (Hamamatsu) and Volocity software (PerkinElmer). Images within each panel were evenly exposed and processed using FiJi (Schindelin *et al.* 2012) and Photoshop CC (Adobe) except for those in Figure 5 where large fluctuations in intensity necessitated different exposure times.

### Western Blot

Log phase yeast were lysed by freezing a pellet, suspending them in Alternate Thorner Buffer (8M Urea, 5% SDS, 40 mM Tris 6.8, 0.1 mM EDTA, 0.4 mg/mL bromophenol blue, 1% β-mercaptoethanol) with glass beads, heating at 70°C for 5 min and then vortexing 1-2 min. An amount of lysate equivalent to 0.1 OD_600_ of yeast was run on an SDS-PAGE gel, transferred to nitrocellulose paper and blotted with mouse anti-HA (Abeam ab18181) followed by donkey anti-mouse conjugated to IR-Dye 800CW (Mandel Scientific 926-32212). The blot was imaged on an Odessey 9120 Infrared Imager (LI-COR).

### Flow Cytometry

Log phase yeast grown in synthetic complete media were measured on an Accuri C6 flow cytometer (BD Biosciences). For each sample 10,000 events were collected and the mean green fluorescence values of the complete ungated populations were recorded.

## Supporting information

Supplementary Information 2 - Recommended protocols

Supplementary Information 1 - Plasmid construction

## ACKNOWLEDGEMENTS

This study was financially supported by a FRQNT Team grant (#208639) to V.J.J.M. and M.W, and a NSERC Discovery (RGPIN/4799) grant to M.W.. B.D.M.B. was supported by a Concordia University Horizon Postdoctoral Fellowship, V.J.J.M. is supported by a Concordia University Senior Research Chair and M.W. is supported by a Canada Research Chair. The authors thank Dr. Elizabeth Conibear at the University of British Columbia and Dr. Jimmy Gollihar at the University of Texas at Austin for sharing strains with parts that were tested for or used to generate components of the toolkit. The authors also acknowledge the Centre for Microscopy and Cellular Imaging funded by Concordia University, Montreal, Canada and the Canada Foundation for Innovation.

## AUTHOR CONTRIBUTIONS

B.D.M.B. designed the research, performed the experiments, and wrote the manuscript with editing from V.J.J.M. and M.W.

## SUPPLEMENTARY DOCUMENTS

**SUPPLEMENTARY FIGURE 1. Additional measures of fluorescence.**

**SUPPLEMENTARY TABLE 1 – MyLO Toolkit Plasmids**

**SUPPLEMENTARY TABLE 2 – Primers used**

**SUPPLEMENTARY TABLE 3 – Yeast strains used**

**SUPPLEMENTARY INFORMATION 1. Plasmid construction details**

**SUPPLEMENTARY INFORMATION 2. MyLo CRISPR Toolkit instructions and recommended protocols**

## Citations in Supplementary Sections

(Bean *et al.* 2018)

## ABBREVIATIONS

CAS: CRISPR-associated protein
CRISPR: Clustered Regularly Interspaced Short Palindromic Repeats
DSB: double-stranded break
ESCRT: endosomal sorting complexes required for transport
GFP: green fluorescent protein
GOI: gene of interest
gRNA: guide RNA
HA: hemagglutinin
HDR: homology directed recombination
NHEJ: non-homologous end joining
OD600: Optical Density at 600 nm
PAM: protospacer adjacent motif
2xPH: tandem repeat of phospholipase Cδ Pleckstrin homology domains
RFP: red fluorescent protein
tSNARE: target soluble N-ethylmaleimide-sensitive factor attachment protein receptor

