## Supplementary Information 2 - Recommended protocols for "The MyLo CRISPR-Cas9 Toolkit: A Markerless Yeast Localization and Overexpression CRISPR-Cas9 Toolkit"

**SUPPLEMENTARY INFORMATION 2: Toolkit details, instructions, and recommended protocols**

The goal of this document is to improve the accessibility of CRISPR/Cas9 with this kit. Therefore, in addition to kit-specific guidelines we include overviews of common CRISPR uses and lab protocols.

**How to choose which Cas9 vector to use**

The MyLO kit contains four Cas9 vector types that differ based on protospacer location, and the selections present:

**(A) No protospacer, complete selection**

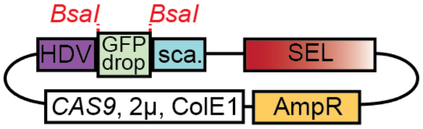

pBBK1, 3, 5, 7, 9

**Uses:**
-Quick, lower efficiency transformations based on gRNA recombination in cell
-Best if only doing a transformation a few times
-Could be used for pre-cloning, but not recommended given Figure 1d/e/f

**(B) No protospacer, split selection**

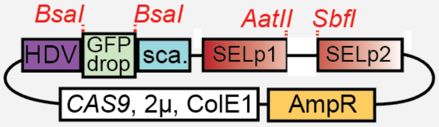

pBBK2, 4, 6, 8, 10

**Uses:**
-Use for pre-cloning guides
-Best for common or low efficiency transformations
-Not for in-cell gRNA recombination

**(C) Pre-cloned protospacer, split selection**

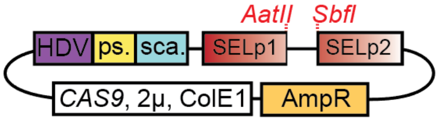

pBBK11-24

**Uses:**
-Use for integration at Sites 1-7 with KanR or HygR selections
-Convenient for integrations at safe harbor loci, otherwise inflexible

**(D) No protospacer, single complete selection**

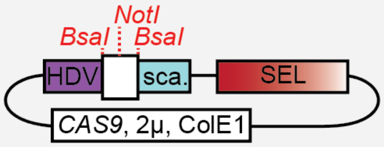

pBBK94, 95

**Uses:**
-Best for quick, lower efficiency transformations based on gRNA recombination in cell
-Similar to first pCAS type, but only contains a single selection (KanR or HygR) for yeast and E. coli making it smaller and more efficient
-Protospacer site has a stuffer with two *BsaI* sites and a *NotI* site improving digestion

In short, for quick transformations use (A) or (D), for higher efficiency or common transformations pre-clone guides into (B) and for integrating into the safe harbor sites presented in this toolkit use (C).

### How to transform with the different types of Cas9 vectors

**(A, ~D) No protospacer, complete selection**

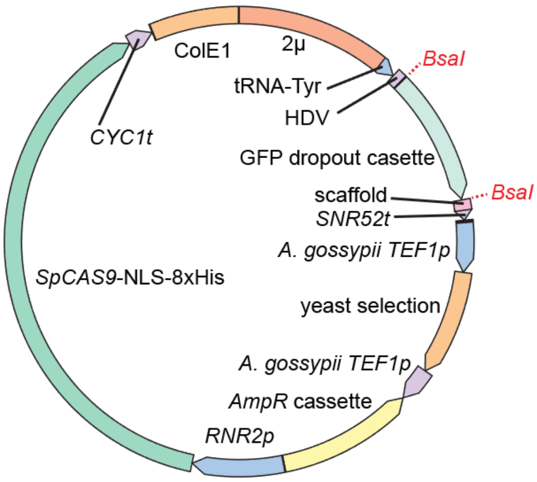

1) Purify vector from *E. coli* grown in media with Ampicillin. *Culture should be green.*
2) Digest 100 ng/µL vector with *BsaI* to release GFP dropout. *Purification is unnecessary.*
3) Design primers encoding a protospacer with 50bp homology to the HDV and the scaffold. Hybridize the primers. *If protospacer is AT-rich, extend primer overlap to increase melting temperature.*

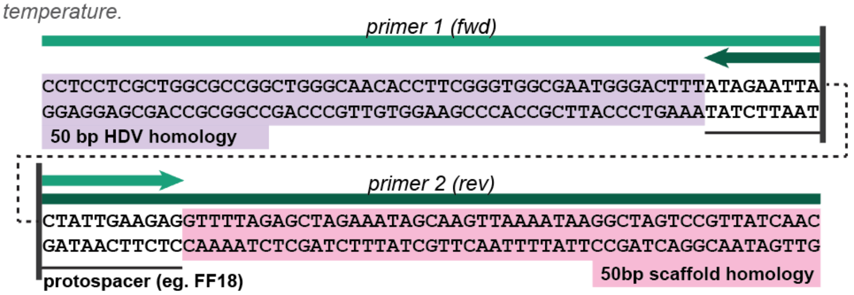

4) Transform yeast with digested pCas9, dimerized primers and donor DNA. A 150 µL Gietz LiAc transformation uses 3-6 pmol dimerized primer, 300-600ng digested pCas9 and 600-800ng of each donor DNA.
**Differences with Type D pCas9: 1) use Kanamycin or Hygromycin B selection, 2) cultures should be white and 3) NotI can be included to improve digest*

**(B) No protospacer, split selection**

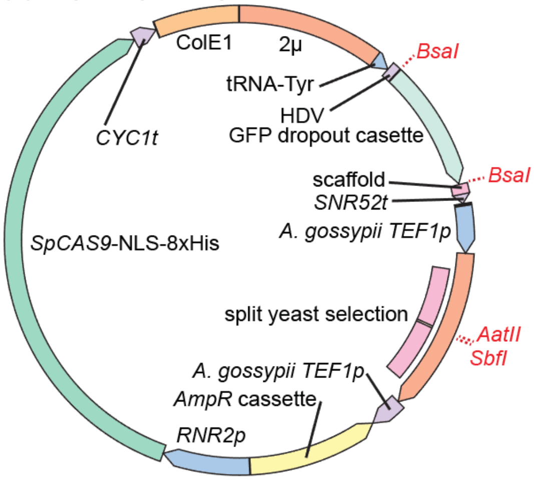

1) Purify pCas9 from *E. coli* grown in media with Ampicillin. *Culture should be green.*
2) Anneal primers designed such that the protospacer led by two thymines are double stranded leaving compatible overhangs for the pCas9. *The thymines restore residues removed for Golden Gate compatibility (CTTT and GTTT overhangs would be too similar).*

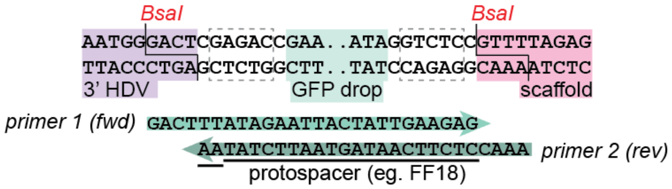

3) Run a *BsaI* Golden Gate reaction with the pCas9 and annealed primers.
4) Transform *E. coli* with reaction, select white colonies, recover pCas9 and optionally confirm by sequencing. *Save this pCas9 for future use.*
5) Digest 100 ng/µL pCas9 with *AatII/SbfI-HF* to linearize. *Purification is unnecessary.*
6) Transform yeast with digested pCas9 and donor DNA. A 150 µL Gietz LiAc transformation uses 300-600ng digested pCas9 and 600-800ng of each donor DNA.

**(C) Pre-cloned, split selection**

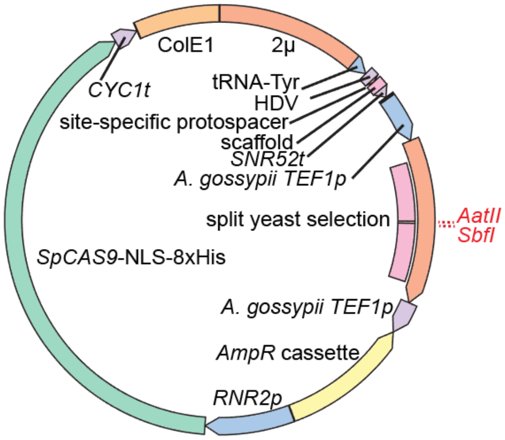

1) Purify pCas9 from *E. coli* grown in media with Ampicillin. *Culture should be white.*
2) Digest 100 ng/µL pCas9 with *AatII/SbfI-HF* to linearize. *Purification is unnecessary.*
3) Transform yeast with digested pCas9 and donor DNA. A 150 µL Gietz LiAc transformation uses 300-600ng digested pCas9 and 600-800ng of each donor DNA.

5' > 3' text of type A/D homology arms (5' > 3'):
**HDV homology,** CCTCCTCGCTGGCGCCGGCTGGGCAACACCTTCGGGTGGCGAATGGGACTTT
**Scaffold homology,** GTTGATAACGGACTAGCCTTATTTTAACTTGCTATTTCTAGCTCTAAAAC

**How to introduce new selectable markers into pCas9**

Recognizing that in some cases additional selectable markers on the pCAS will be useful, the *URA3* pCAS (pBBK09) was designed to simplify exchanging the selectable marker for another complete or split selection by *BsmBI* Golden Gate cloning.

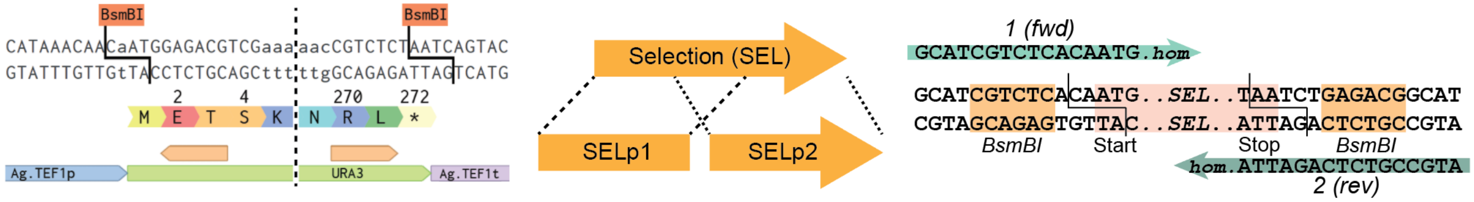

5'>3' text of primer termini: **1,** GCATCGTCTCACAATG; **2,** ATGCCGTCTCAGATTA

The *URA3* gene is flanked with *BsmBI* sites **(left)** that can excise *URA3* by cutting at CAAT and AATC which partially encode START and STOP codons. Choose which type of selection is desired and, if using a split selection, decide where to fragment the selection into two parts. Note that while using roughly the first and last two thirds of the selectable marker works well, expanding the sizes of each part increases the amount of homology for HDR, which may further improve transformation efficiencies **(middle)**. If swapping with a complete selection, amplify the marker with two primers containing homology (hom.) to the 5' and 3' ends of the gene and encode *BsmBI* recognition sites that will introduce CAAT and AATC-compatible overhangs **(right)**. Note that the primers should have 4-6 arbitrary 5' terminal residues to allow efficient *BsmBI* binding. The pCAS and amplified selection can now be used in a two-part *BsmBI* Golden Gate reaction.

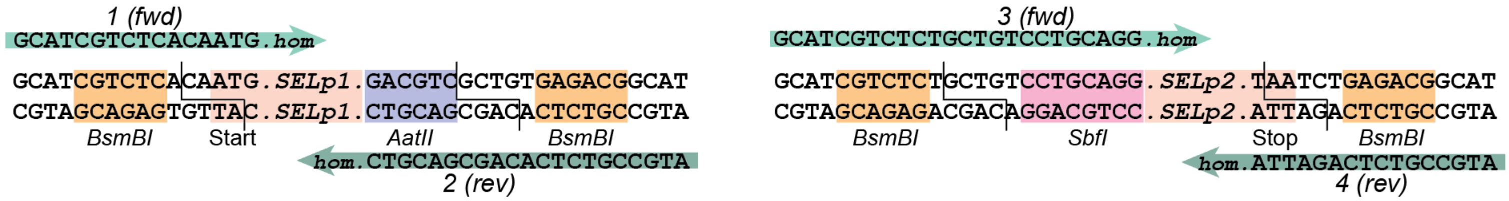

5'>3' text of primer termini: **1,** GCATCGTCTCACAATG; **2,** ATGCCGTCTCACAGCGACGTC; **3,** GCATCGTCTCTGCTGTCCTGCAGG; **4,** ATGCCGTCTCAGATTA

If creating a new split selection cassette, the design of the external primers remains the same but two new internal primers are introduced. The first part of the selection (SELp1) should be amplified with a reverse primer containing an interior *AatII* site and an exterior *BsmBI* site **(left)**. The *BsmBI* overhang is arbitrary but must match that of primer 3 **(right)**. For simplicity GCTG is recommended. For the second part of the selection (SELp2) a new forward primer is used with an internal *SbfI* site and an external *BsmBI* site. Amplifications of SELp1 and SELp2 can be used with pBBK09 in a three-part *BsmBI* Golden Gate reaction to generate the new split selection pCAS.

**Introducing GOIs into integration cassettes, using them, and using cassettes without cloning**

Cloning a gene of interest (GOI) into one of the parent integration vectors (pBBK25-45) differs depending on whether it will be untagged/C-terminal-tagged or N-terminal-tagged because the two cases use different overhangs. In most cases C-terminal tagging is sufficient or preferred so we recommend starting with that vector set and only tagging the N-terminus when there is reason to believe C-terminal tagging would be disruptive. For either vector set, the GOI interest is first amplified to introduce BsaI sites required for prior to Golden Gate cloning with BsaI to introduce it into a parent vector. Note that the primers shown below also contain a *BsmBI* site so that GOIs can optionally be introduced into the entry vector pYTK001 (Lee et al., 2015) using a *BsmBI* Golden Gate reaction and sequenced prior to cloning into the parent vectors as presented here.

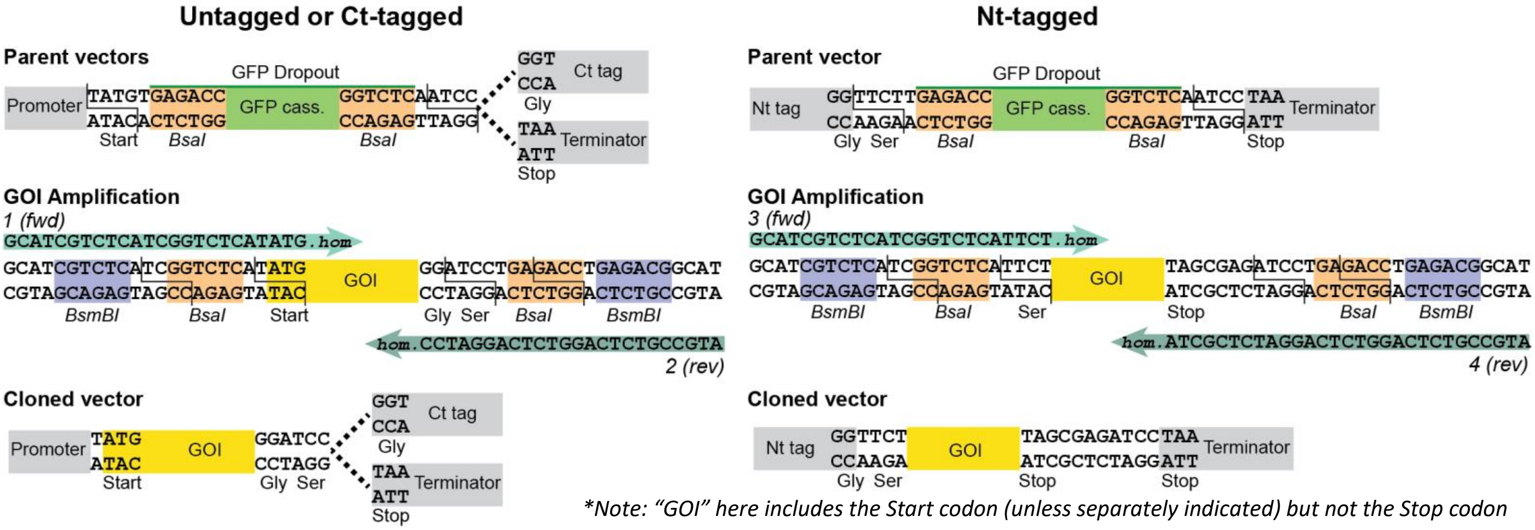

5'>3' text of primer termini: **1,** GCATCGTCTCATCGGTCTCATATG; **2,** ATGCCGTCTCAGGTCTCAGGATCC; **3,** GCATCGTCTCATCGGTCTCATTCT; **4,** ATGCCGTCTCAGGTCTCAGGATCTCGCTA

Both sets of parent vectors contain a GFP dropout cassette that is replaced with the GOI, though the overhangs and flanking regions differ as shown. For untagged or C-terminally tagged GOIs, amplifications are done with a forward primer (1) containing a *BsaI* site that will cut at the Start codon of the GOI and a reverse primer (2) which introduces C-terminal glycine and serine residues at a *BsaI* cut site. These residues contribute to the linker and are necessary for the C-terminal tag or the Stop codon to be in frame. While the two amino acids are small and polar there is a chance that they could be disruptive in the context of the untagged GOI. If this is a concern, reverse primer (4) can be used instead. When amplifying a GOI for N-terminal tagging a forward primer (3) with a BsaI cut site at a serine residue and a longer reverse primer (4) that introduces a Stop codon directly after the GOI are used.

In both cases the amplified GOIs are cloned into an appropriate parent vector in a two-part BsaI Golden Gate reaction.

In some situations, it may be desirable to avoid cloning a GOI into the parent vectors. This may be due to time constraints or an abundance of BsaI sites withing your GOI. To achieve this, digest the target parent vector with *NotI* and *BsaI*, releasing desired fragments with (i) HomL-promoter-(Nt-tag?) and (ii) (Ct-tag?)-terminator-HomR. Co-transforming these two fragments with a pCAS and a GOI amplified using primers with 50bp of homology upstream and downstream of the cloning site should yield positive colonies. Gel purifying the digests will decrease background insertion of the empty cassette and larger amounts of DNA may be required.

**How to make new integration cassettes**

While the MyLO toolkit is limited to integration cassettes with three promoters and three types of tags, the collection can be expanded. As outlined in the plasmid construction supplement, the integration cassettes were built using the modular cloning toolkit structure (Lee et al., 2015), which breaks plasmids down into eight parts with some subparts (see below). To generate new integration cassettes, individual parts can be amplified with primers containing *BsaI* sites that introduce the overhangs indicated below. Templates for these parts could be existing cassettes, the modular cloning toolkit or other sources. It is strongly recommended to read the article on the modular cloning toolkit before generating new integration cassettes.

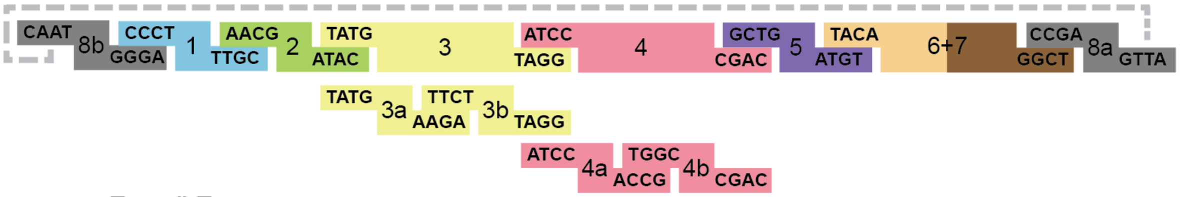

| Type # | Type | Type # | Type | Type # | Type |
|---|---|---|---|---|---|
| 8b | Left Homology Arm | 3a | Nt tag | 4b | Terminator |
| 1 | Left Connector | 3b | GOI/dropout | 5 | Right Connector |
| 2 | Promoter | 4 | Terminator | 6+7 | Right Homology Arm |
| 3 | GOI/dropout | 4a | Ct tag | 8a | Plasmid Backbone |

To facilitate the construction of alternate cassettes, a parent vector containing a GFP dropout replacing parts 2-4 (promoter to terminator) has been provided in the kit (pBBK96). As an example, the simplest case of using this cassette is shown below. Here the GFP dropout is replaced by a new promoter, gene, and terminator. Note that the Start codon for the GOI is in the BsaI overhang (T**ATG**) and that the Stop codon should be included. After the running the Golden Gate reaction, it would be transformed into *E. coli* and white colonies would be selected for confirmation by sequencing.

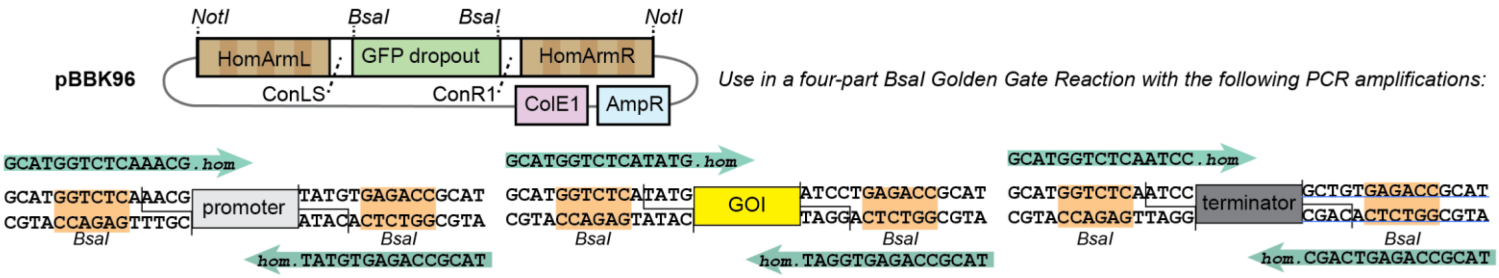

**Overview of strategies for different uses of CRISPR/Cas9**

The primary focus of this toolkit is integrations at specific safe harbor loci. However, the pCAS vectors included can be used for several other markerless manipulations of yeast genomic DNA. Outlined below are the main alternate use cases, which all take advantage of yeast homologous recombination of regions as short as 35-50bp (indicated with dashed Xs).

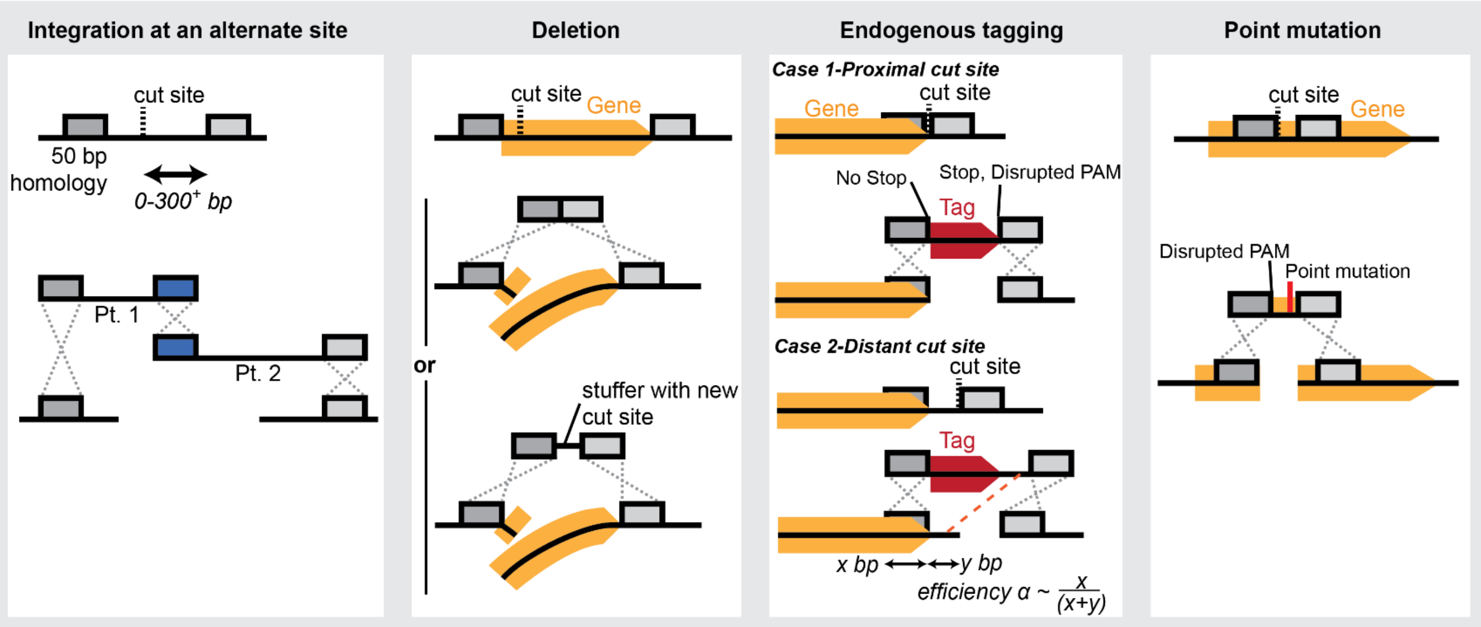

***Integrations at alternate sites***

Within a target locus identify a Cas9 cut site and use a pCAS-gRNA to cut it. Target donor DNA can be introduced in one to eight parts (with efficiency decreasing as part number increases) that contain 50 bp overlapping regions, terminating in homology on either side of the cut site. We recommend to placing the peripheral homology regions 0-300 bp from the cut site, though further also works.

***Deletions***

To delete a gene design and introduce a pCAS-gRNA that cuts within the gene. Also introduce a short donor DNA with homology to the DNA flanking the gene. Optionally the donor can contain an arbitrary new Cas9 cut site to facilitate future manipulations including restoration of the gene. In either case, hybridization of two primers is the easiest way to generate the donor DNA.

***Endogenous Tagging***

In the example of C-terminal tagging, identify a cut site as close to the Stop codon as possible and introduce a pCAS-gRNA to cut there. In an ideal situation with a nearby cut site, amplify the tag with flanking homology to either side of the cut site ensuring that there is no Stop codon preceding the tag, that there is a Stop codon after the tag and that the used cut site is disrupted (which may require mutation of the PAM). In less ideal situations an extended region of the terminator may need to be included, potentially requiring two pieces of donor DNA. Note that in this case recombination can occur in the other terminator region (red dashed line) returning the wild type sequence. The efficiency of achieving the desired integration is proportional to the size of the upstream and internal terminator homology regions.

***Point mutations***

Use a pCAS-gRNA to cut near the site of your desired point mutation and introduce a donor DNA with a silent mutation disrupting the PAM in addition to the desired point mutation. In some cases with distant cut sites it may be necessary to use two donor DNAs.

### Common protocols

Use of this kit requires several common lab protocols. As a resource we are including our protocols, though we would like to point out that many other methods and reagents can be used.

#### *Cas9 cut site selection*

The best cut sites will have a good RNA structure, not have off target sites in the genome, and will have low nucleosome occupancy so the DNA is exposed. To aid in selecting a site based on the first two criteria we recommend using the CCTop server <https://cctop.cos.uni-heidelberg.de:8043/> (Stemmer et al., 2015). Nucleosome occupancy at a target gene can be obtained using a CRISPRi tool <http://lp2.github.io/yeast-crispri/> (Smith et al., 2016).

#### *Restriction digests*

Digests when testing this kit were typically performed with 2x excess enzymes from New England Biolabs (NEB) and 100 ng/µL DNA:

Eg.

| | |
|---|---|
| x µL | 10 µg plasmid |
| 1 µL | Enzyme ( 20 U/µL) |
| 10 µL | 10x Cut Smart Buffer |
| 89-x µL | $dH_2O$ |

*Incubate 2-16 h at 37°C
*Heat inactivate when possible as specified in enzyme documentation (frequently 20 min at 80°C)

#### *Hybridizing Primers*

This refers to annealing and extending a forward and reverse primer with overlapping 3' ends in a short PCR reaction. The only PCR condition that requires changing is the annealing temperature (bold). *Note that the melting temperature in this example is slightly low. Here we would recommend extending the overlap until the melting temperature is ~58-60°C.*

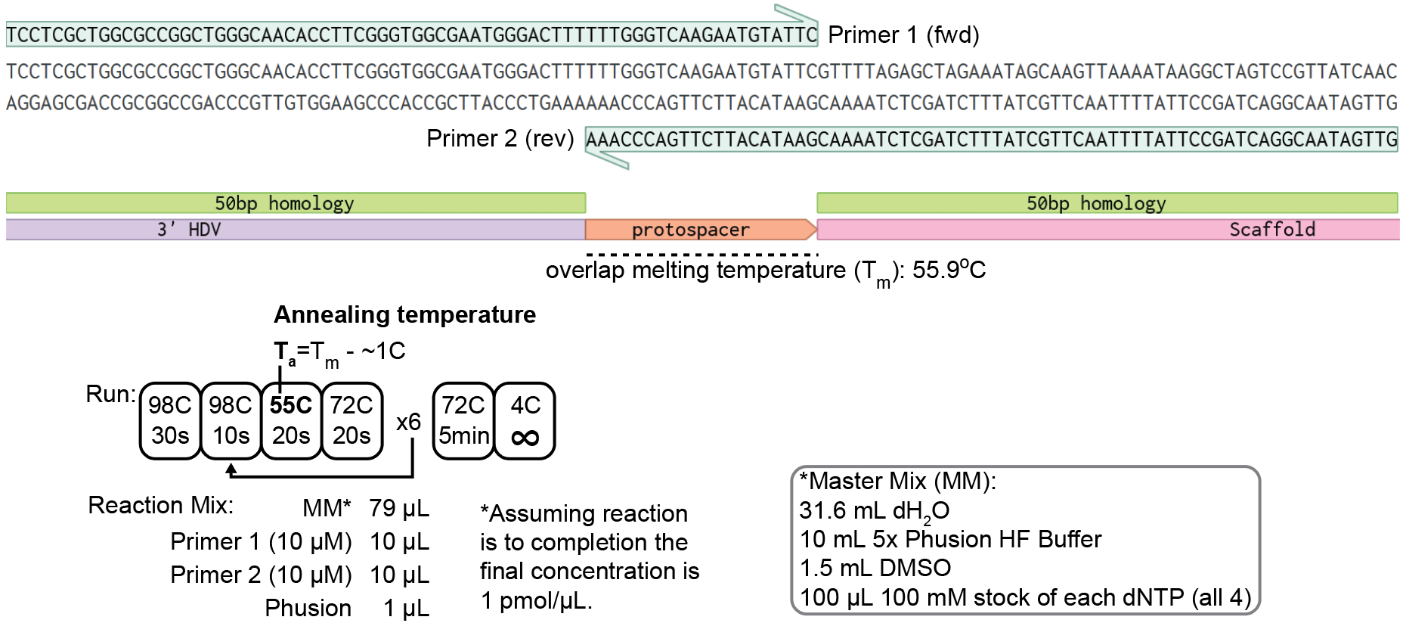

Eg. Hybridizing two primers to introduce a new protospacer into a pCAS by recombination

### *Annealing Primers*

This refers to mixing two primers together, heating them and then slowly dropping the temperature so they anneal while preserving any overhangs

Eg.

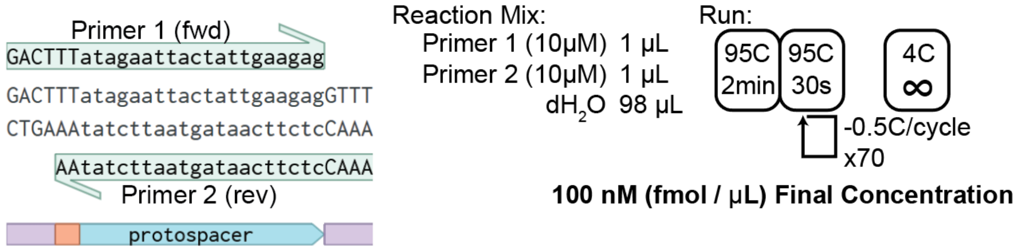

Preparing a protospacer for introduction into a pCAS by Golden Gate cloning

### *Transforming yeast*

We transform yeast using a standard lithium acetate-based protocol (Gietz and Schiestl, 2007) with some adjustments including reduced reaction volumes as follows.

**(1) Prepare yeast:**

-Grow yeast O/N in YPD at 30°C
-Back dilute to 0.3-0.4 $OD_{600}$/mL in YPD or 2xYPD (usually a 1:10-1:15 dilution)
-Grow yeast at 30°C for 4-6 h until 1.2-2.0 $OD_{600}$/mL
-Harvest 3 $OD_{600}$ yeast per transformation reaction
-Wash two times with at least 1 mL $dH_2O$
-Per 3 $OD_{600}$ yeast add:
  100 µL 50% $PEG_{3350}$
  5.6 µL 3 M lithium acetate
  4.4 µL single stranded salmon sperm DNA
    *(prepared by boiling 8 min at 100°C and then freezing; thaw just before use*)
-Mix by vortexing

**(2) Prepare a DNA mix:** *(eg. recombination pCAS transformation)*

-For each reaction:
  6 µL 100 ng/µL BsaI-digested pCAS9
  6 µL 1 pmol/µL hybridized protospacer
  x µL donor DNA (*use ~600-800ng each and attempt to balance the molar ratios*)
  Add $dH_2O$ to a 40 µL final volume

**(3) Transform yeast:**

-Add 110 µL of the yeast preparation to the 40 µL DNA mix (*150 µL final*)
-Vortex to mix
-Incubate 30°C 30min, 42°C 30min
-Spin 4,500 g, remove supernatant
-Two cases:
(A) If auxotrophic selection: resuspend in 500 µL $dH_2O$, plate 150 µL and save remainder at 4°C
(B) If antibiotic resistance selection: resuspend in 500 µL YPD, grow at 30°C for 2.5-16 h, plate 150 µL and save remainder at 4°C

***Golden Gate reactions***

Protocols for Golden Gate reactions depend on the restriction enzyme being used (with $37^{o}C$ or 40-$45^{o}C$ digestion temperatures for *BsaI* and *BsmBI* respectively) as well as the number of parts (with 2-part assemblies efficient in 15 cycles and 8-part assemblies in 25-35 cycles). There are many ways to tweak reactions so we recommend additional resources at NEB <https://international.neb.com/applications/cloning-and-synthetic-biology/dna-assembly-and-cloning/golden-gate-assembly> and the Bennet lab wiki <https://wiki.rice.edu/confluence/display/BIODESIGN/Golden+Gate+Assembly>.

Eg. Cloning a protospacer into a Golden Gate compatible pCAS vector

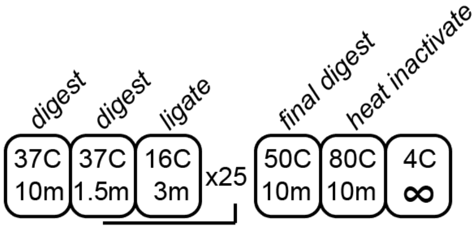

0.5 µL 12.5 fmol/uL pCAS vector
0.5 µL of 100 fmol/uL annealed primers (*2x excess relative to high quality parts as annealing is unlikely complete*)
0.5 µL *BsaI*
0.5 µL T4 ligase
1.5 µL 10x T4 ligase buffer (avoid repeated freeze thaw by using small aliquots)
1.5 µL 10x BSA+PEG (10x: 1 mg/mL BSA, 10% PEG-3350)
10 µL $dH_2O$
/15 µL total rxn
*Store product at $-20^{o}C$ until ready to transform ~7 µL into 25-50 µL chemical competent *E. coli*

***Cleaning internal BsaI/BsmBI sites when Golden Gate cloning***

One roadblock to cloning genes into the integration cassettes (pBBK25-45) is that internal BsaI sites disrupt the Golden Gate reactions. This can be mitigated by ordering the gene with the cut site removed or by skipping cloning and introducing the gene at the site by recombination in the yeast (see paragraph of "Introducing GOIs into integration cassettes, using them and using cassettes without cloning"). Another approach is to amplify the gene in two or more parts with *BsaI*-flanked primers that remove internal *NotI/BsaI* site(s) as shown below. *Note that if genes are initially being cloned into BsmBI-based entry vectors then NotI, BsaI and BsmBI sites must be removed and BsmBI-flanked primers should be used to clean internal sites (as opposed to BsaI-flanked).*

Eg. Designing primers to remove a disruptive *BsaI* site in the interior of a GOI

(1) Identify internal BsaI site to be removed

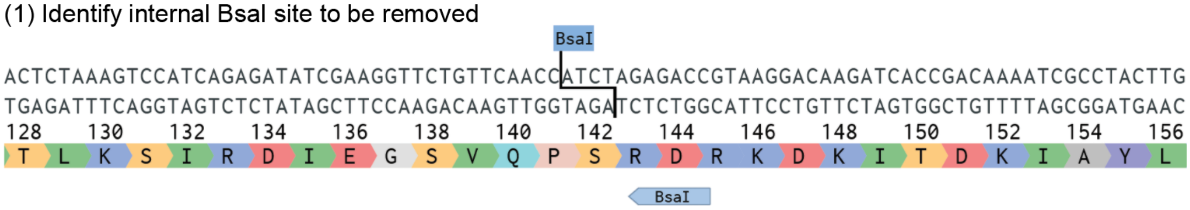

(2) Plan a silent mutation to remove it (here: D GAC>GAT)

(3) Identify a nearby four nucleotide overhang that doesn't interfere with external sites (here: GAGA is compatible with the C-terminal tagging cassette overhangs TATG, TTCT)

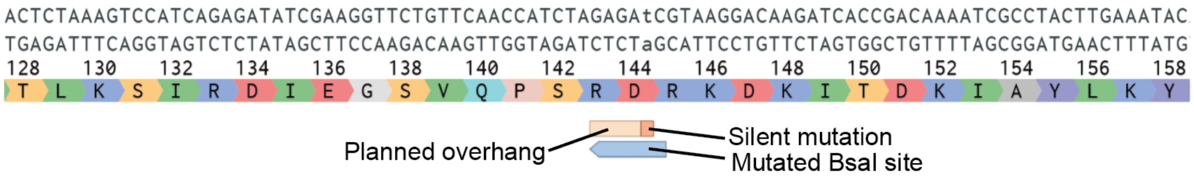

(4) Duplicate the overhang and introduce two nested BsaI sites designed to cut at each overhang sequence

(5) Inside the BsaI sequences add 4-6 arbitrary residues for BsaI binding

(6) Identify the regions outside the overhang sites that will yield good (58 - 60°C) melting temperatures

(7) Annotate each primer: 5' > 3' arbirary residues-BsaI_overhang-homology region

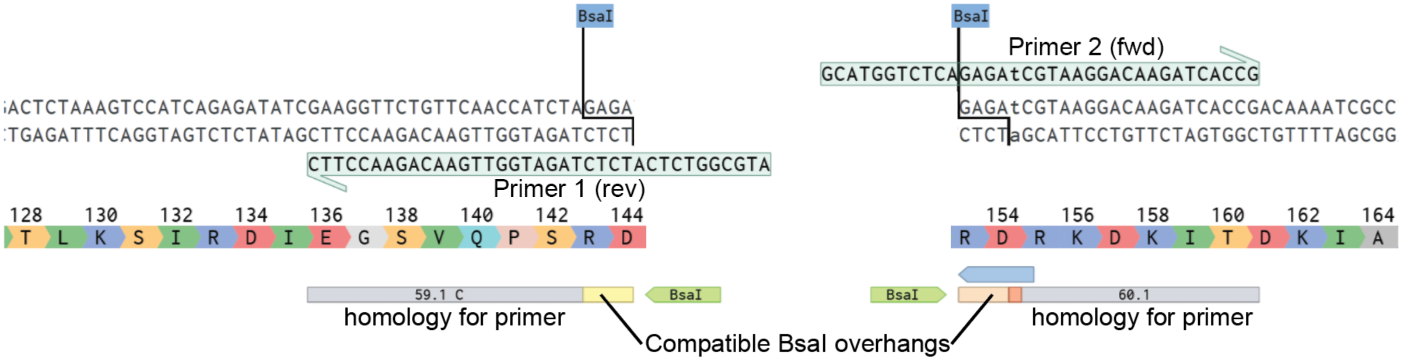
