## Supplementary Figure 1 for "The MyLo CRISPR-Cas9 Toolkit: A Markerless Yeast Localization and Overexpression CRISPR-Cas9 Toolkit"

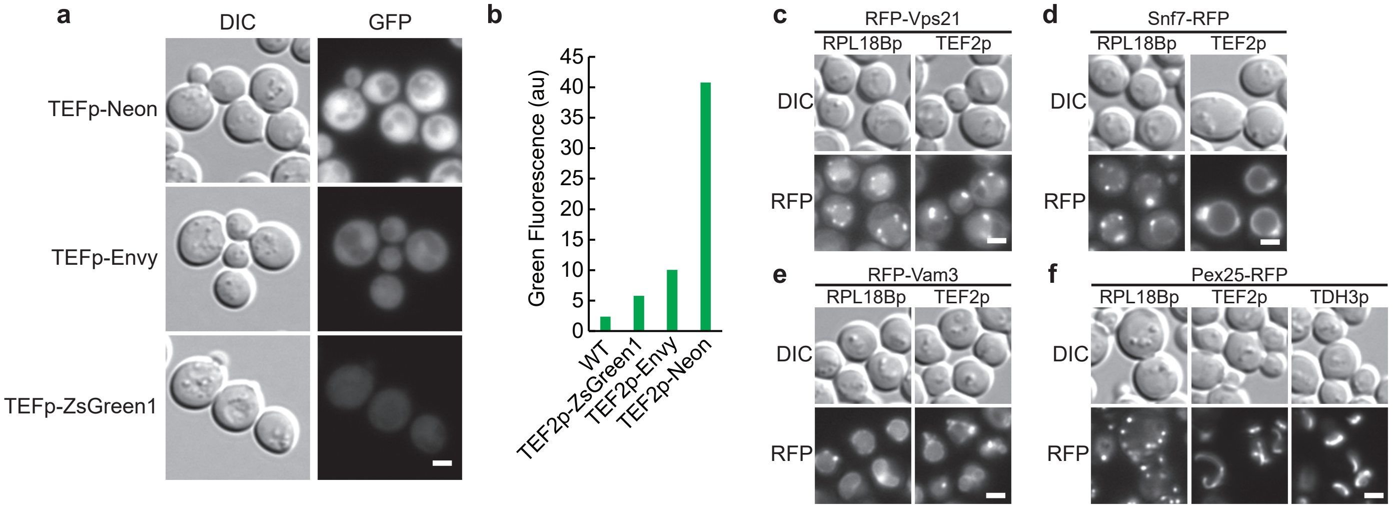


**SUPPLEMENTARY FIGURE 1.** Additional measures of fluorescence. The relative green fluorescence of three tested GFPs expressed from the intermediate *TEF2p* promoter was assessed by **(a)** microscopy and **(b)** flow cytometry. n = 1; 10,000 cells/strain. Fluorescence microscopy of yeast with RFP-tagged markers **(c)** RFP-Vps1, **(d)** Snf7-RFP, **(e)** RFP-Vam3 and **(f)** Pex25-GFP displaying changes in localization as promoter strength increases *RPL18Bp* < *TEF2p* < *TDH3p*. n = 2. Scale bars indicate 2 μm
