## Supplementary Information 1 - Plasmid construction for "The MyLo CRISPR-Cas9 Toolkit: A Markerless Yeast Localization and Overexpression CRISPR-Cas9 Toolkit"

All plasmids were made using either homologous recombination in yeast or by Golden Gate cloning. All vectors were confirmed by sequencing.

### pCAS vectors

Cas9 vectors are derived from pCAS (Addgene #60847) which expresses *S. pyogenes* Cas9 and a *LYP1* gRNA from the tRNA-Tyr RNA polymerase III promoter with a KanR selectable marker (Ryan *et al.*, 2014). The KanR was swapped to HygR to generate pGC1867. The *LYP1* protospacers of the KanR and HygR Cas9 vectors were swapped with a 40bp stuffer containing a *NotI* cut site flanked by two *BsaI* sites to generate pBBK94 and pBBK95 respectively. To do so, a tRNA-Tyr-3'HDV fragment upstream of the protospacers was amplified from pCAS using primers BB29/BB33 and a downstream Scaffold-*SNR52t* fragment was amplified using BB30/34. Both amplicons were gel purified. The two fragments were joined with the 40bp stuffer using splicing by overlap extension (SOE) with BB29/BB30 at full concentration and internal primers encoding the stuffer, BB35/BB36, at $1/125^{th}$ the concentration. The SOE product was purified and co-transformed with *BglI/BglII*-digested pCAS or pGC1867 into yeast.

pBB134 was then constructed from pBBK94 by simultaneously changing the KanR selection for a *BsmBI*-flanked *URA3*, introducing an AmpR *E. coli* selection cassette and removing a *BsmBI* site near the 5' end of *CAS9*. Here pBBK94 was *BsmBI*-digested, the larger fragment was gel purified and co-transformed into yeast with four overlapping PCR products: (i) *URA3* from amplified from pYTK074 (Addgene #65181)(Lee *et al.*, 2015) with BB997/BB998, (ii) *AgTEF1* amplified from pBBK94 using BB999/1000, (iii) an AmpR cassette from pYTK089 (Addgene #65196) amplified with BB1001/1002 and (iv) a fragment containing *RNR2p* and 5' *CAS9* amplified from pBBK94 with BB1003/1004.

pBB134 was converted into the primary toolkit parent, pBBK9, by substituting the 2x*BsaI*+*NotI* protospacer stuffer with a *BsaI*-flanked Golden Gate-compatible GFP dropout. pBB134 was *BsaI*-digested and co transformed into yeast with a GFP dropout fragment amplified from pYTK047 (Addgene #65154) using BB1079/BB1080.

pBBK9 served as the backbone for *BsmBI* Golden Gate reactions to generate Cas9 vectors with different complete and split selectable markers. Complete selections were introduced as single PCR products, while split selections were introduced in two parts as follows. KanR-based selections were amplified from pYTK077 (Addgene #65184) using BB1010/BB1013 for the complete gene in pBBK1, or BB1010/BB1011 and BB1012/BB1013 for the two parts in pBBK2. HygR-based selections were amplified from pYTK079 (Addgene #65186) using BB1105/BB1106 for the complete gene in pBBK3, or BB1105/BB1107 and BB1108/BB1106 for the two parts in pBBK4. NatR-based selections were amplified from pFA6a-natNT2 (Janke *et al.*, 2004) using BB1115/BB1116 for the complete gene in pBBK5, or BB1115/BB1117 and BB1118/BB1116 for the two parts in pBBK6. *LEU2*-based selections were amplified from pYTK075 (Addgene #65182) using BB1110/BB1111 for the complete gene in pBBK7, or BB1110/BB1112 and BB1113/BB1111 for the two parts in pBBK8. The split*URA3* selection parts for pBBK10 were amplified from pYTK074 using BB1120/1122 and BB1123/1121.

pBBK2 (splitKanR) and pBBK4 (splitHygR) were used as backbones in *BsaI* Golden Gate reactions to generate vectors with pre-cloned protospacers. Protospacer inserts were made by annealing primers with Golden gate compatible overhangs: site 1-FF18 BB1125/BB1126, site 2-FF21 BB1127/BB1128, site 3-UserXII-2 BB1129/BB1130, site 4-UserXII-5 BB1131/BB1132, site 5-ARS308 BB1133/BB1134, site 6-ARS416 BB1135/BB1136, site 7-HIS3 BB1137/BB1138. The vectors resulting from the respective introduction of the listed protospacers into pBBK2 were pBBK11-16 and for pBBK4 were pBBK17-24.

**Integration cassette parent vectors**

All vectors containing integration cassettes were assembled using Golden Gate cloning based on a modular cloning toolkit (Lee *et al.*, 2015). Lee *et al.* provide a system by which complete yeast plasmids are assembled from 8 primary part types with compatible Golden Gate overhangs. Most parts are maintained in an entry vector based on pYTK001 (Addgene #65108) and all new parts developed here were first introduced into pYTK001 and sequence confirmed prior to use. We used a slightly altered assembly strategy (Supplementary Materials and Methods Figure 1).

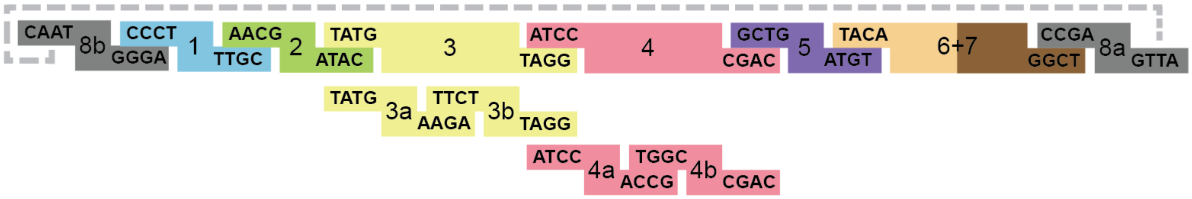


| Type # | Type | Parts of Parent Cassettes (from modular cloning kit, pYTK# + Addgene#, or this work, pBB#) |
|---|---|---|
| 8b | Left Homology Arm | HomL (pBB104) |
| 1 | Left Connector | ConLS (pYTK002, #65109) |
| 2 | Promoter | *RPL18Bp* (pYTK017, #65124), *TEF2p* (pYTK014, #65121), *TDH3p* (pYTK009, #65116) |
| 3 | GOI/dropout | GFPdropout (pBB194) |
| 3a | Nt tag | *Neon(gfp)*-Link (pBB129), *Scarlet(rfp)*-Link (pBB314), *3HA*-Link (pBB215) |
| 3b | GOI/dropout | GFPdropout (pBB195) |
| 4 | Terminator | *SSA1t* (pYTK052, #65159) |
| 4a | Ct tag | Link-*Neon(gfp)* (pBB128), Link-*Scarlet(rfp)* (pBB315), Link-*3HA* (pBB247) |
| 4b | Terminator | *SSA1t* (pYTK062, #65169) |
| 5 | Right Connector | ConR1 (pYTK067, #65174) |
| 6+7 | Right Homology Arm | HomR (pBB103) |
| 8a | Plasmid Backbone | ColE1/*AmpR* (pYTK089, #65196) |

**Supplementary Materials and Methods Figure 1.** The design strategy for the integration cassette parent vectors. Up to nine parts were assembled with the BsaI Golden Gate overhangs indicated. Part types and overhangs are based on the Lee *et al.* modular cloning toolkit with parts 6, 7 and 8b modified as homology arms.

New parts for the integration cassette parents were generated as outlined here prior to using flanking *BsmBI* sites for introduction into pYTK001. The left homology arm in pBB104 was ordered from Twist Biosciences as HomArmL(8b). The right homology arm in pBB103 was assembled into pYTK001 in an eight-part reaction with individual homology region amplifications from BY4741 generated using primer pairs BB731/BB732 (FF18), BB733/BB734 (FF21), BB735/BB736 (UserXII-2), BB737/BB738 (UserXII-5), BB739/BB740 (ARS308), BB741/BB742 (ARS416), BB743/BB744 (HIS3). The GFPdropout cassettes in pBB194 and pBB195 were respectively amplified from pYTK047 using BB759/BB761 and BB760/BB761. Neon(gfp) and Scarlet(rfp) were respectively amplified from a gene order based on pFA6a-ymNeonGreen-CaURA3 (Addgene #125703) and pMD449, a derivative of pFA6a-ymScarletI-CaURA3 (Addgene #118457) provided as a gift from Dr. Elizabeth Conibear, UBC. Neon(gfp)-Link (for pBB129) and Scarlet(rfp)-Link (for pBB314) amplifications were respectively done using BB903/BB904 and BB1180/BB1181. Link-Neon(gfp) (for pBB128) and Link-Scarlet(rfp) (for pBB315) amplifications were respectively done using BB901/BB902 and BB1182/BB1183. 3HA-Link (for pBB215) and Link-3HA (for pBB247) were respectively generated by hybridizing BB987/BB988 and BB1149/1150. The linkers used were GSGPSGPTG (all N-terminal tags), GSGPSGPTG (GFP and RFP C-terminal tags) and GSPSG (3HA C-terminal tag; truncated for cloning reasons).

Parts in sequenced entry vectors were used to make the integration cassette parents (pBBK25-45) via eight- or nine-part end-on-ligation *BsaI* Golden Gate reactions with the components used indicated in the final plasmid descriptions. One intermediate cassette (pBBK96) with a GFPdropout replacing the promoter, GOI, and terminator was similarly made using pBB104, pYTK002, pYTK047, pYTK067, pBB103 and pYTK089.

**Vectors with compartmental marker integration cassettes**

The integration cassette parents were used to make cassettes to express RFP or GFP-tagged compartmental markers, or different types of GFP, as outlined in the results section. Genes encoding compartmental markers were primarily amplified from BY4741 genomic DNA, introduced into pYTK001 using a *BsmBI* Golden Gate reaction and sequenced prior to use. These genes were amplified using primers BB886/BB887 (*RPC19*), BB907/BB908 (*SPC42*), BB1365/BB1366 (*TUB1*), BB896/BB897 (*ARC35*), BB770/BB771 (*SCS2* transmembrane helix), BB953/BB954 (*SED5*), BB959/BB960 (*SFT2)*, BB1355/BB1356 (*VPS21*), BB927/BB928 (*SNF7*), BB950/BB951 (*YPT7*), BB1216/BB1217 (*VAM3*), BB915/BB916 (*PRC1*), BB947/BB948 (*ATG8*), BB932/BB933 (*ERG6*), BB935/BB936 (*PDR16*), BB930/BB931 (*PEX25*) and BB811/BB812 (*TOM70*). In some cases, genes contained an internal *BsaI* or *BsmBI* site which was removed by amplifying the gene in two parts. Here, the internal primers were designed such that they introduced a silent mutation removing the unwanted cut site and contained a compatible *BsmBI* that did not introduce any other mutations into the gene. These two parts, amplified with BB1089/BB1090 and BB1091/BB1092 for *SRM1* or BB1358/BB1359 and BB1360/BB1361 for *PIL1*, were introduced into pYTK001 in three-part Golden Gate reactions.

Additional targets originated from other sources. The *preCOX4* sequence was made by hybridizing BB772/BB773. The tandem *R norvegicus* phospholipase C delta isoform PH domain repeat (2xPH) was amplified in two parts from pBB91 (pRS416-TEF1pr-Link-mCherry-Link-2xPH(PLCd)-ADHt, a gift from Dr. Elizabeth Conibear, UBC) using BB809/BB888 and BB810/BB889 to remove an internal *NotI* cut site. Envy GFP was amplified from pLC2652 (pFA6alink-Envy-SpHis5, a gift from Dr. Elizabeth Conibear, UBC, derived from Slubowski *et al.* (Slubowski *et al.*, 2015)) using BB764/BB1093 and ZsGreen1 GFP was amplified from JG04v2 (a gift from Dr. Jimmy Gollihar, UT Austin) using BB1095/BB1096.
